# Genome-wide, evolutionary, and stress-responsive landscape of the Pectin methylesterase gene family in cucumber and muskmelon

**DOI:** 10.64898/2026.01.08.698443

**Authors:** Anand Kumar Shukla, Bhakti R. Dayama, Shubham S. Nikam, Narendra Y. Kadoo

## Abstract

Pectin methylesterases (PMEs) are key regulators of plant cell wall remodeling; however, their evolutionary dynamics and stress-responsive roles remain poorly understood in cucurbit crops. This study aimed to systematically characterize the PME gene family in cucumber (*Cucumis sativus*) and muskmelon (*Cucumis melo*), addressing how PME diversification, duplication, and regulatory architecture underpin their responses to biotic and abiotic stresses. Using a Hidden Markov Model–based genome-wide screening approach, we identified 52 PME genes in cucumber and 56 in muskmelon, which were classified into Type I and Type II PMEs based on their domain composition. Comparative structural and phylogenetic analyses revealed conserved domain organization but substantial intron-driven structural diversification, resolving PMEs into two major evolutionary lineages with lineage-specific expansion patterns. Duplication and synteny analyses demonstrated that dispersed duplication was the primary driver of PME family expansion, while Ka/Ks estimates indicated strong purifying selection, highlighting functional conservation across cucurbits. Promoter cis-element profiling and protein–protein interaction network analyses revealed extensive enrichment of stress- and hormone-responsive regulatory features, identifying central PME hub genes. Meta-transcriptomic analyses across diverse biotic and abiotic stresses revealed dynamic, condition-specific PME regulation, with Type I PMEs predominantly associated with stress responses in cucumber, whereas both PME types contributed substantially in muskmelon. Several PMEs exhibited conserved stress-induced expression, while others displayed species-, tissue-, or pathogen-specific patterns. Collectively, this study establishes an integrative evolutionary and stress-responsive framework for PME genes in cucurbits, providing mechanistic insights into cell wall plasticity and identifying candidate PME targets for improving multi-stress resilience in crop breeding.

## 1. Introduction

The plant cell wall is a dynamic and multifunctional structure that provides mechanical support to the cell while regulating cell growth and development (Houston et al. 2016; Delmer et al. 2024; Lin et al. 2025). Continuous cell wall remodeling underlies key biological processes, including cell expansion, fruit development and ripening, and defense against pathogens (Haas et al. 2021; Lin et al. 2025; Gallemí et al. 2025). It is also essential for plant adaptation to abiotic stresses through changes in cell wall composition and mechanical properties (Wang et al. 2013; Wu et al. 2018). Pectin is a major polysaccharide of the primary cell wall and middle lamella, where it regulates cell wall porosity, mediates cell-cell adhesion, and maintains structural integrity (Wang et al. 2013; Song 2022; Lin et al. 2025). The functional properties of pectin are primarily determined by the degree of methylesterification of its homogalacturonan (HG) backbone, a linear polymer of galacturonic acid residues (Wang et al. 2013; Wu et al. 2018; Gallemí et al. 2025).

Pectin methylesterases (PMEs) are enzymes that play a crucial role in modifying pectins by catalyzing the removal of methyl ester groups from homogalacturonan, thereby regulating the degree of pectin esterification and its associated physicochemical properties (Wang et al. 2013; Kumar et al. 2023; Lin et al. 2025). Through this activity, PMEs modulate cell wall rigidity and porosity. Controlled, blockwise demethylesterification promotes Ca²⁺-mediated cross-linking and cell wall stiffening (Zhu et al. 2024; Gallemí et al. 2025), whereas excessive or random demethylesterification results in wall loosening, increased porosity, and enhanced susceptibility to pathogens or cell expansion (Lin et al. 2025; Gallemí et al. 2025). Thus, PMEs finely balance cell wall remodeling to coordinate plant growth and defense. The PME gene family is large and structurally diverse in plants (Sun et al. 2022; Song 2022). Based on domain organization, PMEs are classified into two major types: Type I PMEs (proPMEs), which contain an N-terminal pectin methylesterase inhibitor (PMEI)-like domain in addition to the catalytic PME domain, and Type II PMEs, which consist solely of the catalytic domain (Wang et al. 2013; Sun et al. 2022; Song 2022). The PMEI domain confers autoinhibition during protein maturation and transport (Wang et al. 2013; Lin et al. 2025), enabling the precise spatial and temporal regulation of PME activity and underscoring the functional specialization of the gene family (Wang et al. 2013).

PMEs play multifaceted roles in plants and are involved in various growth and developmental processes, including root development, stem elongation, and pollen tube growth (Bosch et al. 2005), as well as fruit ripening and plant stress responses (Jolie et al. 2010; Huang et al. 2022). Moreover, they also play a crucial role in seed germination through seed coat development and breaking of seed dormancy (Ren and Kermode 2000), cell wall stiffening (Siedlecka et al. 2008), defense and stress response (Liu et al. 2018), and controlled cell wall remodeling (Wu et al. 2018). Emerging evidence also confirms the roles of PMEs in abiotic stress tolerance, including responses to salt, cold, drought, heat, and osmotic stress, where they help maintain cell wall integrity (Sun et al. 2022; Cheng et al. 2022; Zhu et al. 2024). In addition to their structural functions, PMEs have been implicated in regulating stomatal movement, thereby contributing to heat stress tolerance through improved transpiration control (Huang et al. 2017; Wu et al. 2017). During plant-pathogen interactions, PME-mediated pectin remodeling influences pathogen penetration (Lionetti et al. 2017; Liu et al. 2018) and can generate damage-associated molecular patterns (DAMPs) that activate immune signaling, highlighting their role at the site of pathogen infection (Lionetti et al. 2017; De Lorenzo and Cervone 2022). Pathogens can thus induce PMEs to alter the plant cell wall and facilitating their entry and spread in the plant (Collmer and Keen 1986). Transient expression of the proPMEs gene in *Nicotiana benthamiana* leaves significantly enhanced tobacco mosaic virus-induced RNA interference (RNAi), demonstrating the effectiveness of PME as an RNAi enhancer. This transitory expression also demonstrated PME’s involvement in transgene-induced gene silencing (Dorokhov et al. 2006).

Genome-wide analyses of PME gene families across diverse plant species have revealed their evolutionary expansion and functional diversification, with 66 PMEs in *Arabidopsis thaliana* (Louvet et al. 2006), 105 in *Linum usitatissimum* (Pinzón-Latorre and Deyholos 2013), 43 in *Oryza sativa* (Jeong et al. 2015), 80 in *Gossypium arboreum* (Li et al. 2016), 43 in *Zea mays* (Zhang et al. 2019), 127 in *Glycine max* (Wang et al. 2021), 66 in *Camellia sinensis* (Huang et al. 2022), 28 in *Diospyros kaki* (Zhang et al. 2022), 21 in *Juglans regia* (Qin et al. 2024), 81 in *Pyrus bretschneideri*, 92 in *Malus domestica*, 62 in *Fragaria vesca*, 65 in *Prunus mume*, 15 in *Pyrus communis*, and 81 in *Pyrus pyrifolia* (Zhang et al. 2024). However, in cucurbit crops such as cucumber (*C. sativus*) and muskmelon (*C. melo*), the functional relevance of PME genes under diverse biological contexts remains incompletely understood. In particular, their potential involvement in plant–pathogen interactions and their differential regulation across multiple biotic and abiotic stress conditions remain largely unresolved. Cucumber and muskmelon are economically and nutritionally important crops cultivated worldwide, and belong to the Cucurbitaceae family (Falade 2021; Pethybridge et al. 2024). Cucumber is particularly significant, ranking as the fourth most cultivated vegetable globally, valued for its nutritional content, including vitamins C, K, and A, as well as minerals, and its hydrating properties (Falade 2021; Mallick 2022). Muskmelon is also highly popular among consumers and represents a relatively high-value crop (Yano et al. 2020; Pethybridge et al. 2024). Despite their agricultural importance, these cucurbits are highly susceptible to a wide array of biotic stresses, including fungal diseases (e.g., powdery mildew, downy mildew, Alternaria leaf spot, Fusarium crown rot), bacterial diseases (e.g., bacterial wilt caused by *Erwinia tracheiphila*, cucurbit yellow vine disease caused by *Serratia marcescens*), and various viral pathogens (Al-Daghari et al. 2021; Pethybridge et al. 2024). They are also severely impacted by abiotic stresses such as drought, salinity, temperature extremes (chilling, heat), and waterlogging, which can lead to significant crop damage and yield losses (Wu et al. 2018; Mondal et al. 2020; Pethybridge et al. 2024). Given that cell wall remodeling enzymes, such as PMEs, are crucial for stress adaptation in plants, understanding their function in cucurbits is essential for enhancing crop resilience and improving productivity (Wu et al. 2018; Lin et al. 2025).

Despite the economic and nutritional importance of cucumber and muskmelon, as well as their susceptibility to diverse biotic and abiotic stresses, a comprehensive understanding of the PME gene family in these cucurbit crops remains limited. Although PMEs have been characterized in several plant species, existing studies have primarily focused on tissue-specific expression patterns, such as those in roots, flowers, anthers, and pollen, under normal developmental conditions. However, their systematic involvement in biotic and abiotic stress responses has remained largely unexplored. To bridge this gap, the present computational study provides a comprehensive genome-wide and comparative analysis of the PME gene family in cucumber and muskmelon, integrating evolutionary, regulatory, and stress-responsive expression analyses. By combining phylogenetics, duplication and selection pressure analyses, promoter architecture, protein-protein interaction networks, and extensive meta-transcriptomic profiling under diverse stress conditions, this work uncovers both conserved and species-specific PME regulatory strategies. These findings offer new insights into PME-mediated cell wall remodeling during stress adaptation and identify key candidate genes for improving stress resilience in cucurbit crops.

## 2. Materials and methods

### 2.1 Sequence retrieval and identification of PME gene family members

The genome sequence of cucumber (*C. sativus*, Chinese Long v3) (Li et al. 2019) and muskmelon (*C. melo*, DHL92 v4) (Castanera et al. 2020) were downloaded from the Cucurbit Genomics Database (CuGenDBv2; http://www.cucurbitgenomics.org/v2/, Accessed: December 1, 2024) (Yu et al. 2023). To identify PME gene family members, 66 *Arabidopsis thaliana* PMEs (The Arabidopsis Information Resource; https://www.arabidopsis.org/) and 43 PMEs from *Oryza sativa* (Rice Genome Annotation Project; https://rice.uga.edu/) were used as reference datasets. These well-annotated dicot and monocot PMEs were used as queries in homology-based BLASTP searches against the cucumber and muskmelon genomes, enabling the reliable detection of conserved PME homologs across the cucurbit species with e-value ≥ 1e^-5^, identity ≥ 40, and match length ≥ 100 (Song 2022). Following this, a sequence search was conducted using HMMER (version 3.3) with an e-value threshold of 1e^-10^. The search was performed against the protein datasets of both species using Hidden Markov Model (HMM) profiles of the conserved PME (PF01095) and PMEI (PF04043) domains, downloaded from the Pfam database (https://www.ebi.ac.uk/interpro/entry/pfam/#table). To validate the HMM search results, all candidate PME sequences were subjected to domain composition analysis using the NCBI Batch CD-Search tool (Wang et al. 2022) with default parameters against the Pfam database. Only proteins containing the conserved “pectinesterase” domain were retained, and redundant entries were removed. The final non-redundant set of confirmed candidates was designated as members of the PME gene family in cucumber and muskmelon. The corresponding genomic DNA sequences, protein sequences, and coding sequences (CDS) of the confirmed PME genes were retrieved from CuGenDBv2. The identified PME genes were systematically named according to their physical positions on chromosomes, with numbering progressing sequentially from chromosome 1 to the last chromosome in each genome (e.g., CsaPME1–CsaPME52 and CmePME1–CmePME56).

### 2.2 Physicochemical characterization and chromosomal localization of PME genes

The cucumber (*C. sativus*) and muskmelon (*C. melo*) PMEs were designated as CsaPMEs and CmePMEs, respectively. The identified CsaPMEs and CmePMEs proteins were analyzed for their physicochemical properties using the ProtParam tool from ExPASy (http://web.expasy.org/protparam/) (Gasteiger et al. 2005). The analysis included determining the number of amino acids, molecular weight (MW), theoretical isoelectric point (pI), grand average of hydropathicity (GRAVY), aliphatic index, and instability index, all performed with default settings of the ProtParam tool. Additionally, the gene loci, start and end positions, chromosome numbers, and strand information for the identified PME family genes were extracted from CuGenDBv2.

### 2.3 Analysis of gene structure and conserved motif recognition

TBtools (Chen et al. 2020) was used to analyze the gene structures of the identified PME family genes utilizing the GTF file (Hu et al. 2015). Exon–intron organization was visualized, and intron content was calculated to assess structural variation among the identified PMEs. Intron length was defined as the total length of all intronic regions within a gene. Intron content (%) was determined as the proportion of the cumulative intron length relative to the total gene length (including both exons and introns), calculated using: intron content (%) = (total intron length of a gene / total gene length) × 100. Additionally, Multiple Expectation-Maximization for Motif Elicitation (MEME) (version 5.5.7) (https://meme-suite.org/meme/) was used to identify conserved motifs in the pectinesterase protein sequences of *C. sativus* and *C. melo*. The MEME analysis was performed using the classical mode with default settings, specifying 10 as the maximum number of motifs (Bailey and Elkan 1994). Ten different MEME motifs (Motifs 1 to 10) were identified in each of *C. sativus* and *C. melo*. These motifs were tested for enrichment with default e-value using SEA (simple enrichment analysis) (version 5.5.7) (Bailey and Grant 2021). The identified MEME motifs were functionally annotated using InterProScan (version 5.76-107.0; https://www.ebi.ac.uk/interpro/search/sequence/) (Blum et al. 2025; Paysan-Lafosse et al. 2025) to determine their associated protein domains and families.

### 2.4 Phylogenetic analysis

Phylogenetic analysis was conducted to investigate the evolutionary relationships among PME proteins in cucumber (52 CsaPMEs), muskmelon (56 CmePMEs), *O. sativa* (43 PME proteins), and *A. thaliana* (66 PME proteins). A total of 217 PME protein sequences were aligned using MUSCLE (version 5.3) (Edgar 2022). IQ-TREE (version 2.3.6) (Minh et al. 2020) was then used to determine the optimal protein substitution model among the 224 available models using ModelFinder. The best-fit model was selected based on the Akaike Information Criterion (AIC), and a maximum likelihood (ML) phylogenetic tree was constructed with 1,000 bootstrap replicates using the ultrafast bootstrap approximation method. The resulting phylogenetic tree was visualized using iTOL (version 7.2.2) (Letunic and Bork 2024).

### 2.5 *Cis*-regulatory element analysis of PME gene promoters

To predict *cis*-acting regulatory elements in the promoter regions of the PME genes, a 2.0 kb upstream DNA sequence from the transcription start site of CsaPMEs (cucumber) and CmePMEs (muskmelon) was extracted using the Basic Bio Sequence View tool in TBtools. These sequences were then analyzed using the PlantCARE database (http://bioinformatics.psb.ugent.be/webtools/plantcare/html/) to identify known *cis*-regulatory elements (Lescot et al. 2002). The identified elements were categorized into four functional groups: (i) growth and development-related elements, (ii) hormone-responsive elements, (iii) light-responsive elements, and (iv) stress-responsive elements. The results were visualized in R using the pheatmap, RColorBrewer, viridis, grid, and gridExtra packages (Neuwirth 2014; Auguie and Antonov 2017; Kolde 2018; Garnier et al. 2024).

### 2.6 Protein-protein interaction network construction and identification of hub PMEs

The protein-protein interaction (PPI) network was constructed using the STRING database (Szklarczyk et al. 2025) to explore functional associations among proteins. Functional enrichment analysis was performed to identify biological pathways and processes associated with the network. The PPI network was subsequently analyzed to predict hub genes using CytoHubba (Chin et al. 2014), which implements 12 different algorithms for hub gene identification in Cytoscape (Shannon et al. 2003). The resultant scores from each algorithm were normalized to a range of 0 to 1 using the min-max normalization method. An aggregated score was then calculated by integrating the normalized scores from all algorithms, allowing for a comprehensive ranking of PME proteins.

### 2.7 Gene duplication patterns of the PME family

To understand the evolutionary dynamics of PME genes, gene duplication events were analyzed in cucumber (*C. sativus*), muskmelon (*C. melo*), *A. thaliana*, and rice (*O. sativa*). Initially, a BLASTp search was conducted across the whole proteome of each species to identify syntenic gene blocks, using an e-value threshold of 1e^-10^ and a maximum of five hits. The Duplicate Gene Classifier tool in MCScanX (version 1.0.0) (Multiple Collinearity Scan toolkit X) (Wang et al. 2012) was employed to categorize duplicated genes based on their origin.

### 2.8 Synteny analysis

Synteny analysis was performed to investigate the evolutionary relationships among *C. sativus*, *C. melo*, *A. thaliana*, and *O. sativa*. The genome assembly and annotation files (GFF) for cucumber and muskmelon were obtained from CuGenDBv2, while the corresponding files for *A. thaliana* and *O. sativa* were retrieved from Phytozome (https://phytozome-next.jgi.doe.gov) and the Rice Genome Annotation Project (https://rice.uga.edu), respectively. The identification of syntenic blocks was performed using MCScanX with default parameters. The resulting collinear blocks were visualized through dual synteny plots, with PME genes in cucumber and muskmelon specifically highlighted to assess their conservation across species.

### 2.9 Identification of selection pressure acting on the PME gene family

To investigate the selection pressure acting on the PME gene family, we employed OrthoFinder (version 3.0) (Emms and Kelly 2019) to identify one-to-one orthologs of cucumber and muskmelon PME genes across cucurbit crops: watermelon (*Citrullus lanatus*), *Cucurbita maxima* ‘Rimu’, bitter gourd (*Momordica charantia*), bottle gourd (*Lagenaria siceraria*), sponge gourd (*Luffa cylindrica*), wax gourd (*Benincasa hispida*), chayote (*Sechium edule*), monk fruit (*Siraitia grosvenorii*), and snake gourd (*Trichosanthes cucumerina*) available on CuGenDBv2 (Yu et al. 2023). We also identified orthologs of the 52 cucumber PMEs within the 56 muskmelon PMEs, and *vice versa*. The CDS and protein sequences from all species were downloaded, and OrthoFinder was run using default parameters on the protein sequences, retaining only one-to-one orthologous pairs for analysis. Protein sequences of the orthologs were aligned using CLUSTALW (Thompson et al. 2003), and the resulting alignments were processed with PAL2NAL (Suyama et al. 2006) to obtain codon-aware nucleotide alignments of the corresponding CDS. These codon alignments were then analyzed using KaKs Calculator 2.0, applying the universal genetic code and the γ-MYN model (Wang et al. 2009) to calculate the nonsynonymous (Ka) to synonymous (Ks) substitution ratios (Ka/Ks). Selection pressure was categorized based on Ka/Ks values, where Ka/Ks > 1 indicated positive selection, Ka/Ks < 1 suggested purifying (negative) selection, and Ka/Ks ≈ 1 indicated neutral evolution. The distribution of Ka/Ks values for PME orthologs in cucumber and muskmelon was visualized using boxplots to compare evolutionary dynamics within the gene family.

### 2.10 Expression profiling of PME genes under abiotic and biotic stresses

To investigate the role of PME genes in stress responses, their expression profiles were analyzed under various abiotic and biotic stresses in cucumber and muskmelon using publicly available transcriptomic datasets. In cucumber, PME gene expression was examined under 10 abiotic stress conditions, including chilling and cold stress, drought, salt stress (with and without silicon treatment), H₂S-regulated salt stress, long-term waterlogging, waterlogging stress in hypocotyls, and high-temperature stress. Similarly, for biotic stress conditions, PME gene expression was analyzed in response to various fungal, bacterial, and viral infections, including *Podosphaera xanthii* (powdery mildew), *Pseudoperonospora cubensis* (downy mildew), *Pseudomonas syringae* (angular leaf spot), *Alternaria cucumerina* (Alternaria leaf spot), *Fusarium oxysporum* (Fusarium wilt), cucumber green mottle mosaic virus (CGMMV), Prunus necrotic ringspot virus, and nematode. In muskmelon, PME gene expression was examined under various abiotic stress conditions, including cold and chilling stress, salt stress, and waterlogging stress (Arora et al. 2017). For biotic stress, the expression of PME genes was analyzed in response to *Phytophthora capsici*, *Stagonosporopsis cucurbitacearum*, *F. oxysporum*, *P. xanthii* (powdery mildew), and in response to tomato leaf curl New Delhi virus (ToLCNDV).

Transcriptomic data for cucumber and muskmelon were retrieved from CuGenDBv2, and average FPKM (Fragments Per Kilobase of exon per Million reads mapped) values were extracted for both biotic and abiotic stress conditions. The FPKM values from different datasets were merged using an in-house developed Python script, followed by normalization using log₂ transformation with the following formula:

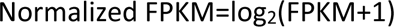

The normalized expression values were then visualized using pheatmap, grid, and gridExtra packages ‘R’ packages to illustrate the differential expression patterns of PME genes under various stress conditions.

To assess structural conservation among the stress-responsive PMEs, representative *Arabidopsis* and rice PMEs, as well as their orthologs in cucumber and muskmelon, were modeled using SWISS-MODEL (Waterhouse et al. 2018). The predicted structures were aligned and superimposed using PyMOL (Schrodinger 2023), and root mean square deviation (RMSD) values were calculated to quantify three-dimensional similarity among the orthologous PME proteins.

## 3. Results

### 3.1 Genome-wide identification and classification of PME genes in cucumber and muskmelon

To systematically identify PME family members, the reference genome sequences of cucumber and muskmelon were screened using HMM-based searches. The identified genes were categorized into two structural classes: Type I PMEs (proPMEs), which contain both the N-terminal PMEI (PF04043) domain and the C-terminal pectinesterase catalytic domain (PF01095); and Type II PMEs, which possess only the catalytic PME domain (PF01095) and lack the PMEI domain. The 52 PME genes identified in cucumber were designated as CsaPMEs, while the 56 PME genes detected in muskmelon were named as CmePMEs **(Table S1)**. Domain composition analysis confirmed that all retained candidates possessed the conserved PME catalytic domain, with variability in the presence of the PMEI domain. Among CsaPMEs, 30 were assigned to Type I **(Figure S1)** and 22 to Type II **(Figure S2)**, while muskmelon comprised 31 Type I **(Figure S3)** and 25 Type II **(Figure S4)** members. The domain structures of cucumber and muskmelon PME proteins are illustrated in **Figures S5** and **S6**, respectively.

### 3.2 Physicochemical characteristics and chromosomal locations of PME proteins

Across the two cucurbits, PME proteins displayed high variation in sequence length, molecular weight (MW), and isoelectric point (pI). For example, in cucumber, the lengths of CsaPMEs ranged from 94 to 605 amino acids (∼10-66 kDa) with pI values of 4.77-9.99, whereas in muskmelon, CmePMEs ranged from 64 to 654 amino acids (∼7-73 kDa) with pI values of 4.95-9.86. For both species, GRAVY scores were predominantly negative (cucumber: −0.345 to −0.005; muskmelon: −0.69 to 0.056), indicating overall hydrophilicity. Aliphatic indices also spanned a broad range (cucumber: 66.09–92.29; muskmelon: 44.92–93.83), and most proteins were predicted to be stable (instability index < 40 for 46/52 in cucumber and 50/56 in muskmelon). Overall, the physicochemical properties of PME were broadly conserved between cucumber and muskmelon, despite variation at the individual gene level **(Tables S1** and **S2)**.

Chromosomal mapping showed an uneven distribution of PME genes in both species. In cucumber, the 52 PME genes were distributed across all seven chromosomes, with the highest density on chromosome 3 (13 genes), followed by chromosome 5 (11 genes), while chromosomes 4 and 6 contained only four genes each **(Figure S7)**. Similarly, the 56 muskmelon PME genes were distributed across all 12 chromosomes, with chromosomes 6 and 9 harboring the largest number (10 genes each). In contrast, chromosomes 7 and 10 contained two genes each, and chromosome 3 contained only a single PME gene **(Figure S8)**.

### 3.3 Gene structure diversity and conserved motif architecture of cucurbit PMEs

Gene structure analysis revealed substantial variability in intron content among the PME genes in both cucumber and muskmelon, underscoring marked structural heterogeneity within the gene family. Type I PMEs generally exhibited a simpler gene structure, comprising 2–3 exons, while Type II PMEs showed increased structural complexity, containing 2–4 exons in cucumber and 4–5 exons in muskmelon. In cucumber, intron content varied widely, ranging from 0.35% in *CsaPME30* to 90.39% in *CsaPME43*, the latter being classified as a Type II PME. Notably, most cucumber PME genes were distributed within intron proportion intervals of 20-40% (16 genes) and 60-80% (16 genes). A similarly broad distribution of intron content was observed in muskmelon as well. The lowest intron proportion was detected in *CmePME29* (0.46%), followed by *CmePME35* (0.51%), whereas the highest intron content was recorded in *CmePME44* (81.53%), belonging to the Type II PME subclass. The majority of muskmelon PME genes also fell within intron ranges of 20-40% (18 genes) and 60-80% (16 genes). This demonstrates extensive intron length variation among the cucumber and muskmelon PME genes, suggesting dynamic evolution of gene structure that may contribute to functional divergence and regulatory complexity within this multigene family **(Tables S3**-**S4; Figures S1-S4)**.

MEME analysis identified ten conserved motifs (Motifs 1-10) across PME proteins of both *C. sativus* and *C. melo*, and subsequent SEA confirmed their significant enrichment, indicating strong conservation and functional relevance of these motifs within the PME gene family. In cucumber, the SEA ranking showed that Motifs 7, 5, 6, and 2 were the most significantly enriched **(Table S5)**. Several of these motifs contain conserved glycine, aspartate, and catalytic residues characteristic of pectin methylesterase catalytic cores. Similarly, in muskmelon, Motifs 5, 2, 4, and 7 were most highly enriched, largely overlapping with those identified in cucumber **(Table S6)**. The near-identical consensus sequences and ranking patterns between the two species indicate a high degree of motif conservation. Importantly, Motif 1, shared between cucumber and muskmelon, represents the core PME catalytic motif, confirming that the fundamental enzymatic mechanism is conserved across species **(Figures S9** and **S10)**.

InterProScan annotation of the MEME-identified motifs revealed that the majority correspond to the conserved pectinesterase catalytic domain (Pfam PF01095; PANTHER PTHR31707) and the pectin lyase-like superfamily (SSF51126) in both cucumber and muskmelon, confirming their functional relevance as bona fide PMEs. Notably, motif 10 in both species was annotated as a plant invertase/pectin methylesterase inhibitor (PMEI) domain (Pfam PF04043), supporting the presence of Type I PMEs with regulatory N-terminal regions. These results validate the accuracy of motif identification and highlight the conserved domain architecture of PME proteins in cucurbits **(Tables S7** and **S8)**.

### 3.4 Phylogenetic relationships and evolutionary grouping of PME family members

Phylogenetic analysis was performed using 217 PME protein sequences from cucumber (52 CsaPMEs), muskmelon (56 CmePMEs), rice (43 OsPMEs), and *Arabidopsis* (66 AtPMEs). Multiple sequence alignment generated 3,291 aligned positions, comprising 2,093 conserved sites, 746 parsimony-informative sites, and 452 singleton sites, indicating substantial sequence conservation alongside evolutionary divergence within the PME family. ModelFinder identified WAG+I+G4 as the best-fit amino acid substitution model based on the Akaike Information Criterion. The phylogeny analysis resolved the PME proteins into two major clusters (**Figure 1**), largely corresponding to their domain architectures. One cluster predominantly consisted of Type II PMEs, while the second cluster comprised Type I PMEs (proPMEs). To further refine the evolutionary relationships, the Type II PME cluster was subdivided into six subclades (Clades I-a to I-f). Likewise, the Type I PME cluster was separated into eight subclades (Clades II-a to II-h), reflecting lineage-specific diversification within each group. Notably, *CmePME29*, although structurally classified as a Type II PME, clustered within the Type I PME group (Clade II-h), suggesting possible evolutionary divergence or domain architecture variation. Additionally, five rice PMEs and one *Arabidopsis* PME (AT5G20860; *PMEI*-*PME54*) did not cluster with any defined subclade, indicating potential species-specific evolutionary trajectories or highly divergent PME members. PMEI-PME54 encodes the pectin methylesterase inhibitor, involved in regulating cell wall modifications.

**Figure 1:**
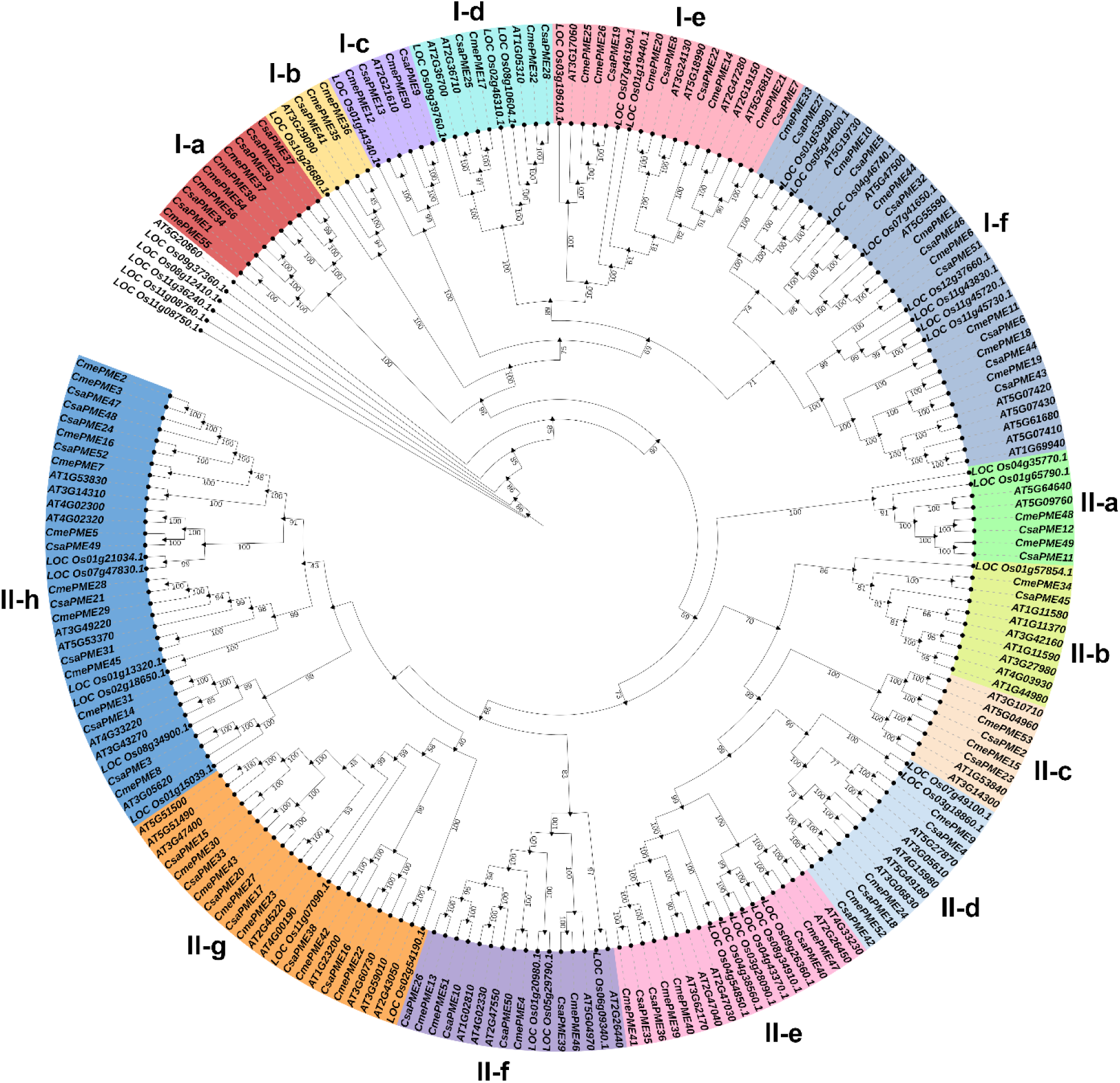
Maximum likelihood (ML) phylogenetic tree depicting the evolutionary relationships among pectin methylesterase (PME) proteins from cucumber (*Cucumis sativus*, 52CsaPMEs), muskmelon (*Cucumis melo*, 56 CmePMEs), rice (*Oryza sativa*, 43 PMEs), and *Arabidopsis thaliana* (66 PMEs). The phylogeny resolves two major clades: Clade I (subclades Ia–If) representing Type II PME proteins, and Clade II (subclades IIa–IIh) representing Type I PME proteins, reflecting their distinct evolutionary origins and domain architectures. Bootstrap support values are indicated at the nodes.

### 3.5 *Cis*-regulatory landscape of PME gene promoters

Promoter analysis revealed that PME genes in both cucumber and muskmelon harbor a diverse array of *cis*-acting regulatory elements associated with light response, phytohormone response, stress responses, and growth and development, indicating complex transcriptional regulation of this gene family (**Figure 2**). In cucumber, light-responsive elements were abundantly represented across nearly all CsaPMEs, with several genes such as *CsaPME1*, *CsaPME6*, *CsaPME23*, and *CsaPME33* containing more than ten light-responsive motifs. Phytohormone-responsive elements, including those related to abscisic acid, methyl jasmonate, auxin, ethylene, salicylic acid, and gibberellin signaling, were widely distributed, with genes such as *CsaPME4*, *CsaPME6,* and *CsaPME19* exhibiting particularly high enrichment (**Figure 2A**). Stress-responsive *cis*-elements constituted a significant fraction of the promoter architecture, with several genes (e.g., *CsaPME5*, *CsaPME45*, and *CsaPME47*) harboring more than 20 stress-responsive motifs. These motifs were predominantly enriched for drought-responsive MYC elements and anaerobic induction-associated ARE motifs, as well as temperature- and pathogen-responsive elements. In contrast, growth- and development-related elements were comparatively fewer and unevenly distributed, although genes such as *CsaPME23* showed notable enrichment **(Table S9)**.

**Figure 2:**
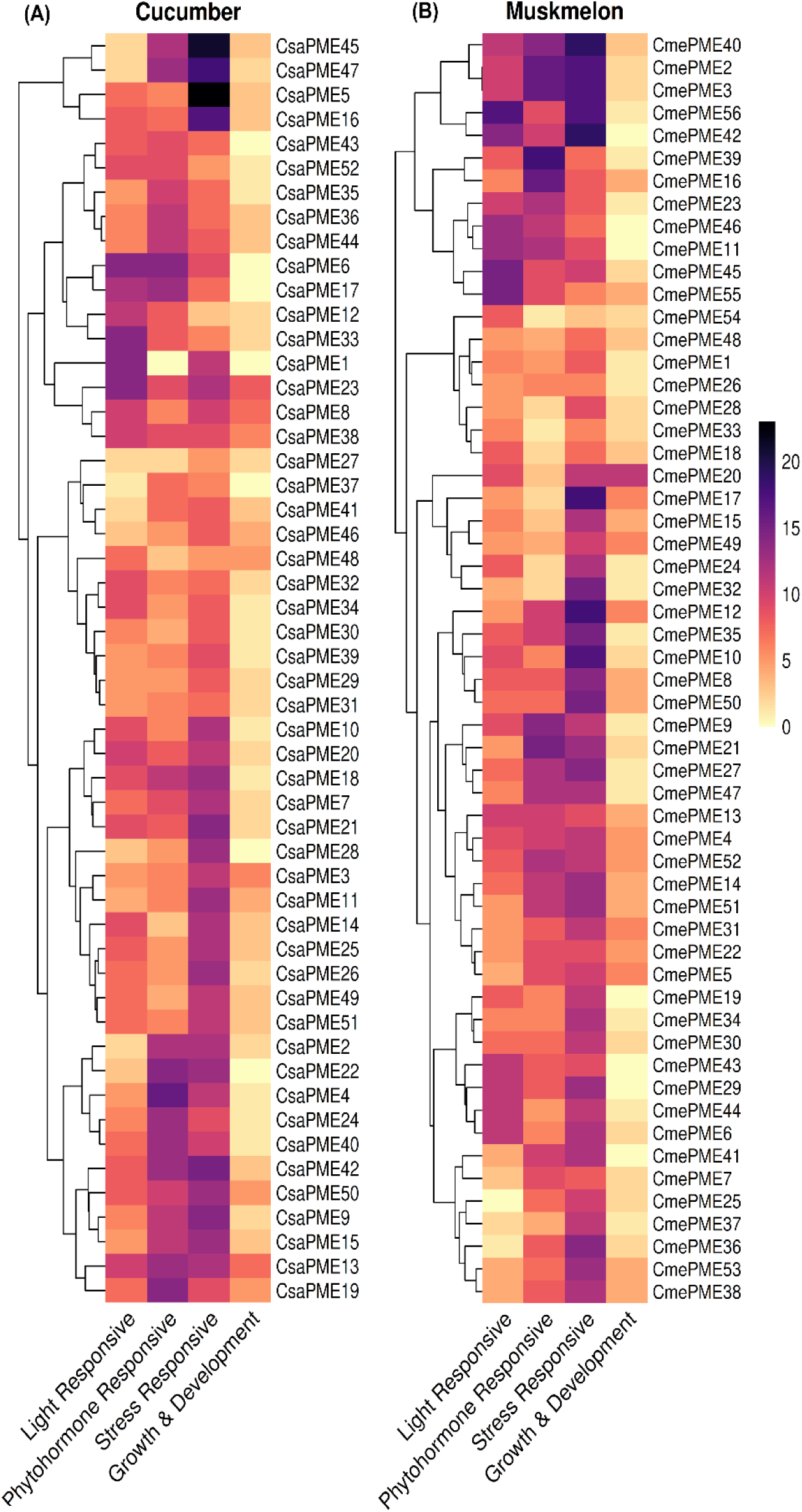
Cis-regulatory element composition in the promoter regions of PME genes. (A) Heatmap depicting the abundance and distribution of cis-acting regulatory elements in the 2.0 kb upstream promoter regions of cucumber (*Cucumis sativus*) PME genes. (B) Heatmap depicting the abundance and distribution of cis-acting regulatory elements in the promoter regions of muskmelon (*Cucumis melo*) PME genes. The identified elements were classified into four functional categories: light-responsive, phytohormone-responsive, stress-responsive, and growth and development-related elements. The heatmaps were generated using the R packages pheatmap, RColorBrewer, viridis, grid, and gridExtra.

Similarly, muskmelon PME promoters displayed extensive *cis*-element diversity (**Figure 2B**). Light-responsive elements were highly prevalent, with genes such as *CmePME56*, *CmePME45*, and *CmePME55* showing strong enrichment. Phytohormone-responsive elements were particularly abundant in several genes, including *CmePME39*, *CmePME2*, *CmePME3*, and *CmePME16*, suggesting hormonal regulation of PME expression. Stress-responsive *cis*-elements were prominently represented across the PME gene family, with several genes (e.g., *CmePME40*, *CmePME42*, *CmePME17*, and *CmePME12*) harboring more than 17 stress-responsive sites, predominantly enriched for drought-responsive MYC motifs. Despite their overall lower abundance, growth- and development-related *cis*-elements were strongly enriched in *CmePME20*, particularly the seed-specific AAGAA motif, indicating possible involvement in seed development **(Table S10)**.

### 3.6 Protein-protein interaction networks reveal central hub PMEs

The PPI network analysis revealed extensive functional associations among PME proteins and other cell wall-modifying enzymes in both cucumber and muskmelon **(Figure S11)**. In cucumber, the PME-centered network comprised 55 nodes and 199 edges, indicating a highly interconnected interaction landscape showing interactions with multiple pectin-degrading enzymes, including pectate lyases (Csa_6G513500, Csa_5G622520, Csa_2G326460) and a glycosyl hydrolase family 28 protein (Csa_3G629740), highlighting a coordinated involvement in pectin remodeling **(Figure S11a)**. CytoHubba analysis identified *CsaPME43*, *CsaPME30*, *CsaPME44*, *CsaPME27*, and *CsaPME46* as the top-ranked hub genes, suggesting their central regulatory roles within the PME interaction network **(Table S11)**. Functional enrichment analysis of the cucumber PPI network revealed a substantial overrepresentation of the pectin catabolic process and cell wall modification among biological processes, accompanied by significant enrichment of pectinesterase, aspartyl esterase, pectinesterase inhibitor, and pectate lyase activities under molecular function categories. Most interacting proteins were predominantly localized to the extracellular region, consistent with PME-mediated cell wall dynamics. A minor but significant enrichment was also observed for defense response to Gram-negative bacteria, suggesting potential roles in stress-associated cell wall remodeling **(Figure S12)**.

In muskmelon, the PME interaction network consisted of 49 nodes and 131 edges, reflecting a slightly less dense but functionally conserved network architecture **(Figure S11b)**. PME proteins interacted primarily with polygalacturonase-like proteins (A0A5A7T6B3, A0A5D3DS41, MPG, MPG2) and pectate lyase (A0A5D3CBM7), reinforcing the coordinated action of multiple pectin-modifying enzymes. Hub gene analysis identified *CmePME19*, *CmePME52*, *CmePME50*, *CmePME9*, and *CmePME13* as the top-ranking central nodes **(Table S12)**. Consistent with cucumber, enrichment analysis in muskmelon highlighted pectin catabolic process and cell wall modification as dominant biological processes, along with significant enrichment of pectinesterase, aspartyl esterase, and enzyme inhibitor activities. The majority of proteins were enriched in the extracellular region, underscoring the conserved extracellular functional niche of PMEs across both species **(Figure S13)**.

### 3.7 Duplication-driven expansion of the PME gene family

Gene duplication analysis revealed distinct evolutionary patterns of PME gene expansion across the four plant species examined: cucumber, muskmelon, *Arabidopsis thaliana*, and rice. In all species, dispersed duplication represented the predominant mode of PME gene expansion, although its relative contribution varied considerably among lineages. In *A. thaliana*, PME genes were mainly derived from dispersed duplications (50%), followed by a substantial contribution from whole-genome/segmental duplications (31.82%), reflecting the impact of ancient polyploidy events in shaping the PME gene family **(Table S13)**. Proximal duplications accounted for 16.67%, while tandem duplication contributed minimally (1.52%). In *O. sativa*, PME expansion was driven by dispersed duplications (79.07%), with a notable contribution from tandem duplications (16.28%). In contrast, whole genome duplication (WGD)/segmental duplication events were not detected, indicating that local and dispersed duplication mechanisms primarily underlie PME diversification in rice. For *C. sativus*, PME genes appear to be predominantly produced through dispersed duplication (75%), followed by tandem duplication (11.54%) and a modest contribution from WGD/segmental duplication (9.62%) events. Proximal duplications were relatively rare (3.85%), suggesting limited local gene expansion. In *C. melo*, dispersed duplication also dominated (64.29%), but a comparatively higher proportion of proximal duplications (25%) was observed, distinguishing muskmelon from the other species. Tandem (3.57%) and WGD/segmental duplications (7.14%) contributed to a lesser extent **(Table S14; Figure 3**).

**Figure 3:**
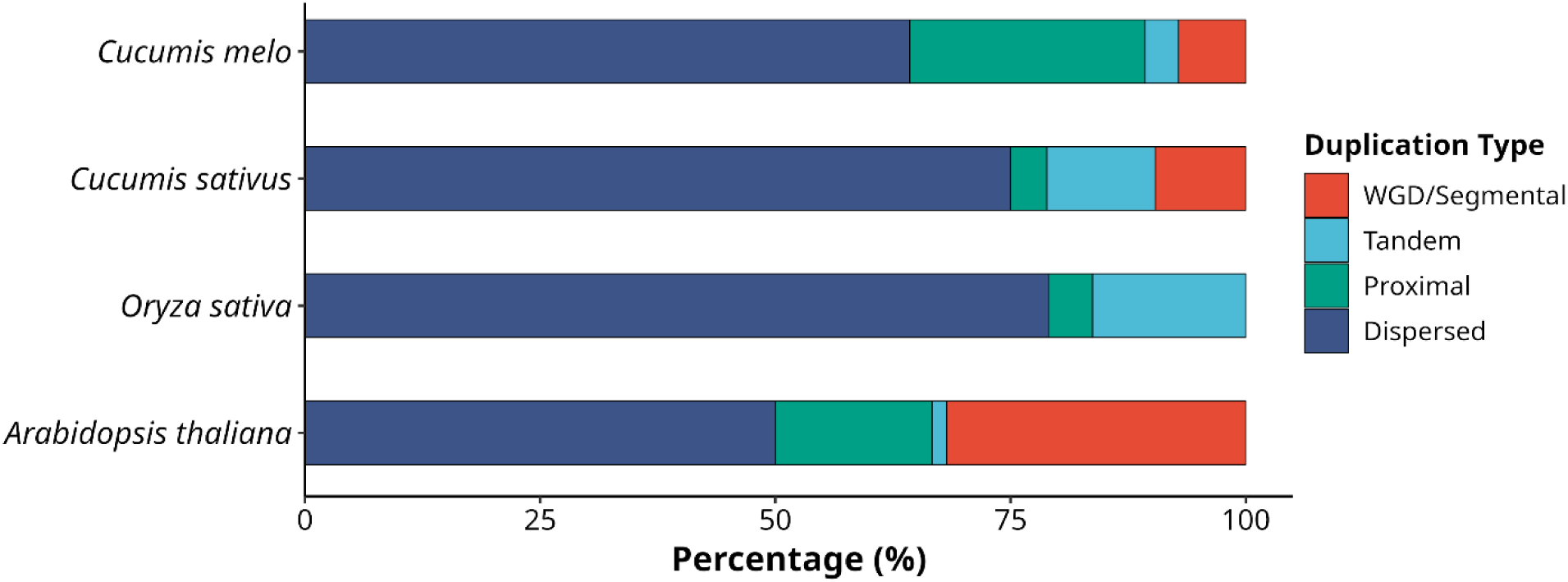
Duplication-driven expansion of the PME gene family across four plant species. Gene duplication patterns of PME genes in cucumber, muskmelon, *Arabidopsis thaliana*, and rice. Genome-wide synteny analysis was performed using BLASTp followed by MCScanX to identify duplicated gene pairs and syntenic blocks. PME genes were classified into different duplication categories—whole-genome/segmental duplication (WGD/SD), tandem, proximal, and dispersed—using the Duplicate Gene Classifier.

### 3.8 Comparative synteny and evolutionary conservation of PME genes across species

Synteny analysis revealed varying degrees of collinearity of PME genes among *C. sativus*, *C. melo*, *A. thaliana*, and *O. sativa*, reflecting their evolutionary relatedness and divergence patterns. A substantial number of conserved syntenic PME gene pairs were detected between the two cucurbit species, with 46 orthologous pairs identified between cucumber and muskmelon, indicating strong genomic conservation and a shared evolutionary origin of PME genes within the Cucumis lineage. A comparative analysis with *A. thaliana* revealed moderate conservation, with 33 syntenic PME gene pairs identified between *Arabidopsis* and cucumber, and an equivalent 33 pairs between *Arabidopsis* and muskmelon. This level of conservation suggests that a significant fraction of PME genes predates the divergence of cucurbits and *Arabidopsis* and has been retained through speciation. In contrast, fewer syntenic relationships were observed with *O. sativa*, consistent with its more distant evolutionary relationship as a monocot species. Only 10 syntenic PME gene pairs were identified between rice and cucumber, and seven pairs between rice and muskmelon, indicating extensive genome rearrangements and lineage-specific gene loss or diversification following monocot-dicot divergence **(Table S15; Figure 4**).

**Figure 4:**
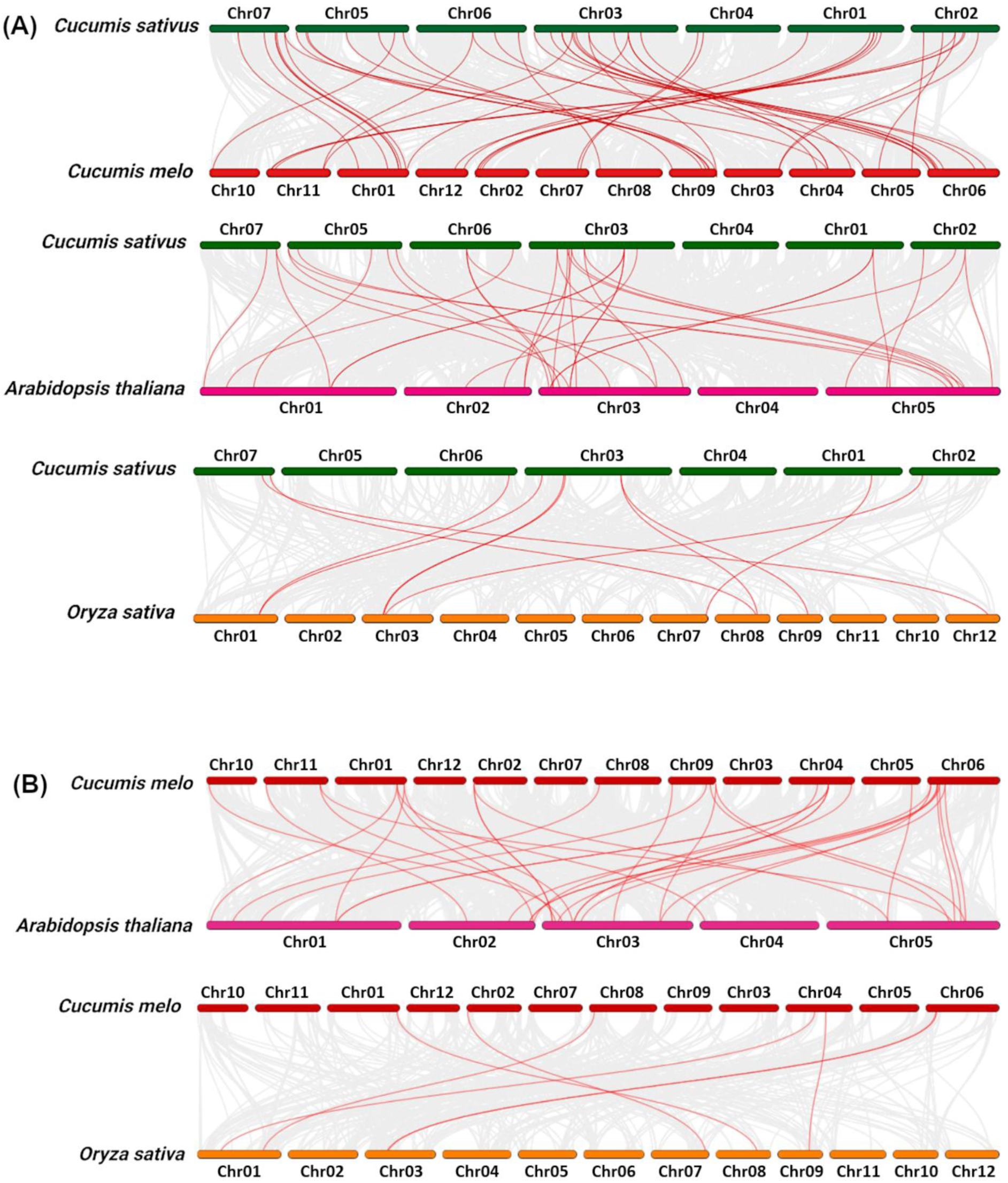
Synteny analysis of PME genes between (A) cucumber and other plant species, and (B) muskmelon and other plant species

### 3.9 Purifying selection shapes the evolutionary trajectory of PME genes

A total of 49 one-to-one orthologous PME genes were identified for cucumber and muskmelon across multiple cucurbit species, including watermelon, *Cucurbita maxima* ‘Rimu’, bitter gourd, bottle gourd, sponge gourd, wax gourd, chayote, monk fruit, and snake gourd. Ka/Ks analysis of these conserved orthologs revealed that, in cucumber, Ka/Ks values ranged from 0.0179 to 0.7860 (**Figure 5**). In contrast, the values ranged from 0.0176 to 0.8283 in muskmelon, reflecting a similar extent of evolutionary divergence across cucurbit lineages. Notably, all orthologous PME genes exhibited Ka/Ks ratios below 1, indicating that the PME gene family is predominantly under strong purifying selection across cucurbit crops. No evidence of pervasive positive selection was observed, suggesting a high level of evolutionary constraint and functional conservation of PME genes within the Cucurbitaceae.

**Figure 5:**
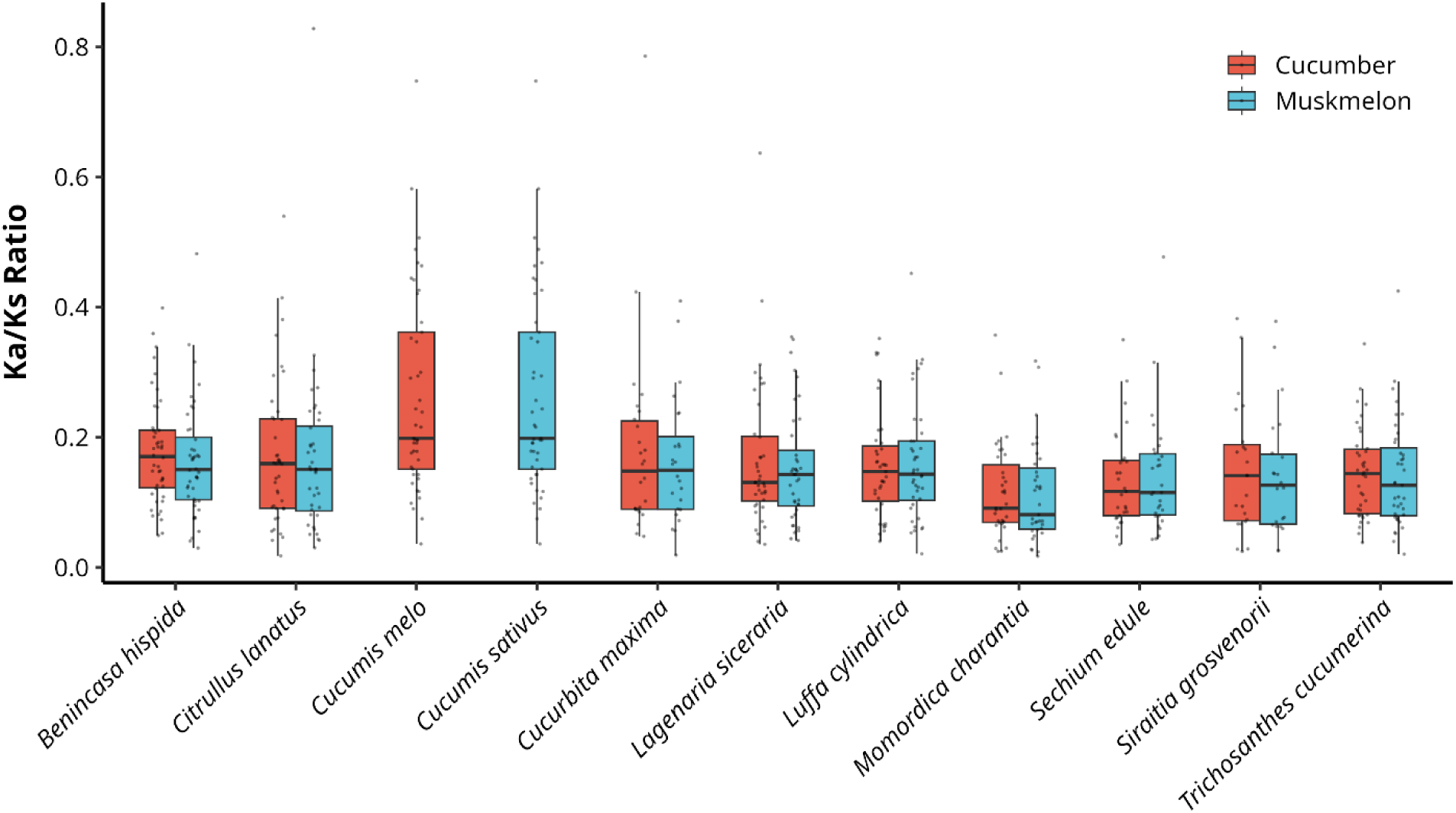
Distribution of Ka/Ks ratios for PME orthologs across cucurbit species. Boxplots show the distribution of nonsynonymous to synonymous substitution rate ratios (Ka/Ks) for one-to-one orthologous PME gene pairs in cucumber (*Cucumis sativus*) and muskmelon (*Cucumis melo*) across cucurbit crops. The boxplots were generated using the R packages tidyverse and ggsci to compare evolutionary constraints acting on PME genes in cucumber and muskmelon. Ka/Ks values were used to infer selection pressure, where Ka/Ks < 1 indicates purifying selection, Ka/Ks ≈ 1 indicates neutral evolution, and Ka/Ks > 1 indicates positive selection. The top and bottom edges of the boxplots indicate the upper and lower quartiles, respectively; the lines within the boxplots indicate medians; the vertical lines above and below the boxplots indicate upper and lower extremes, respectively, while the points above or below the vertical lines indicate outliers.

### 3.10 Stress-responsive expression patterns of PME genes in response to abiotic and biotic stresses

Based on the normalized expression threshold (log₂FPKM ≥ 5), 21 out of 52 cucumber PME genes exhibited variable expression patterns across multiple abiotic and biotic stresses. Among these, 14 belonged to Type I PMEs, whereas seven were classified as Type II PMEs, indicating a predominant contribution of Type I PMEs to stress-responsive expression. Under abiotic stress conditions, 13 Type I and six Type II PMEs were highly expressed, while 12 Type I and four Type II PMEs showed elevated expression under biotic stress. Notably, 11 Type I and three Type II PMEs were commonly induced under both stress categories (**Figures 7** and **8**).

**Figure 6:**
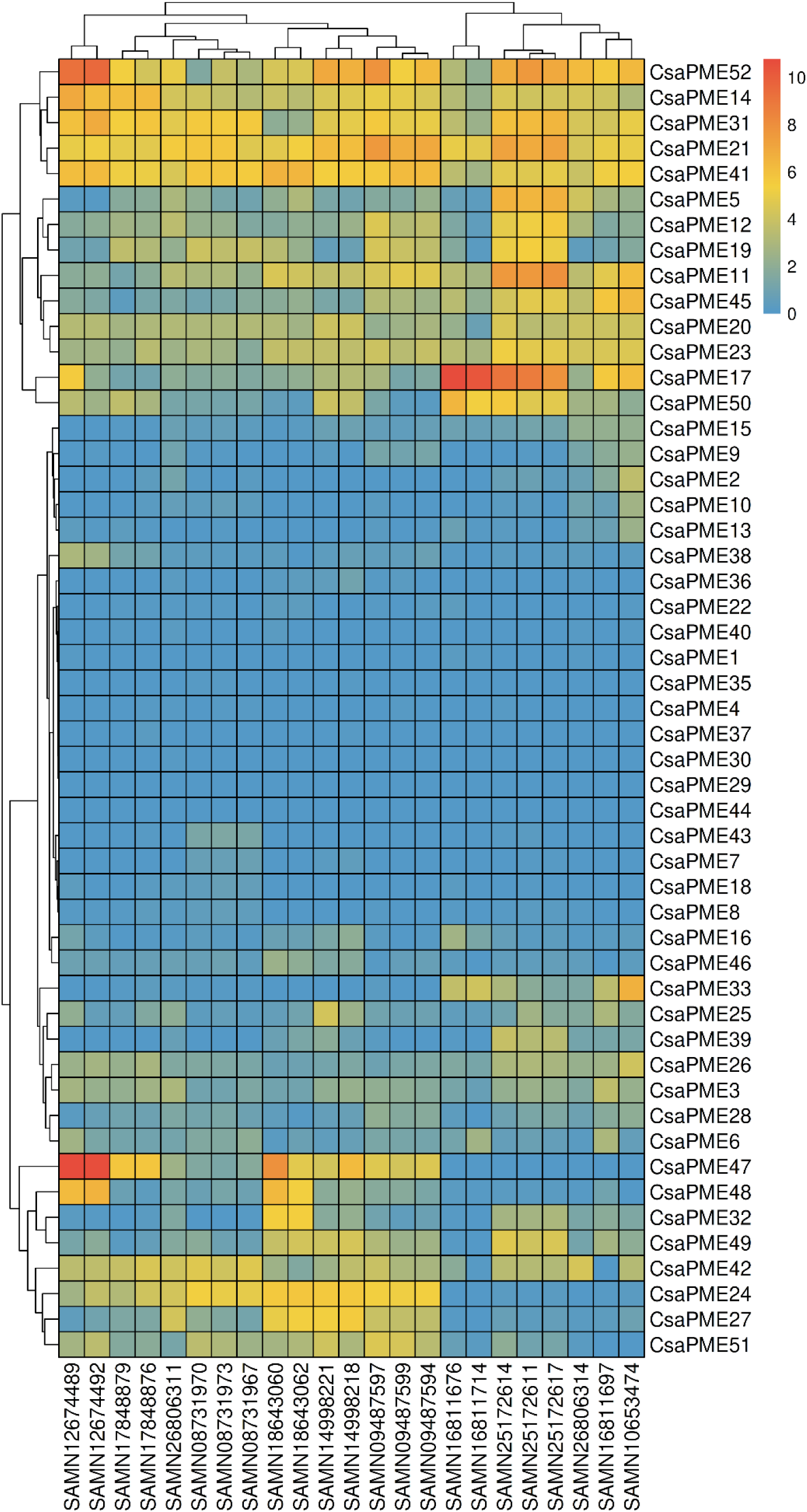
Normalized expression profiles of cucumber PME genes across multiple abiotic stresses. Expression data were retrieved from publicly available RNA-seq datasets, and heatmaps were generated using R packages pheatmap, grid, and gridExtra. Please refer to Table S16 for sample identifiers and the corresponding BioSample accessions and experimental conditions. The color intensity represents relative PME gene expression levels across abiotic stress treatments.

**Figure 7:**
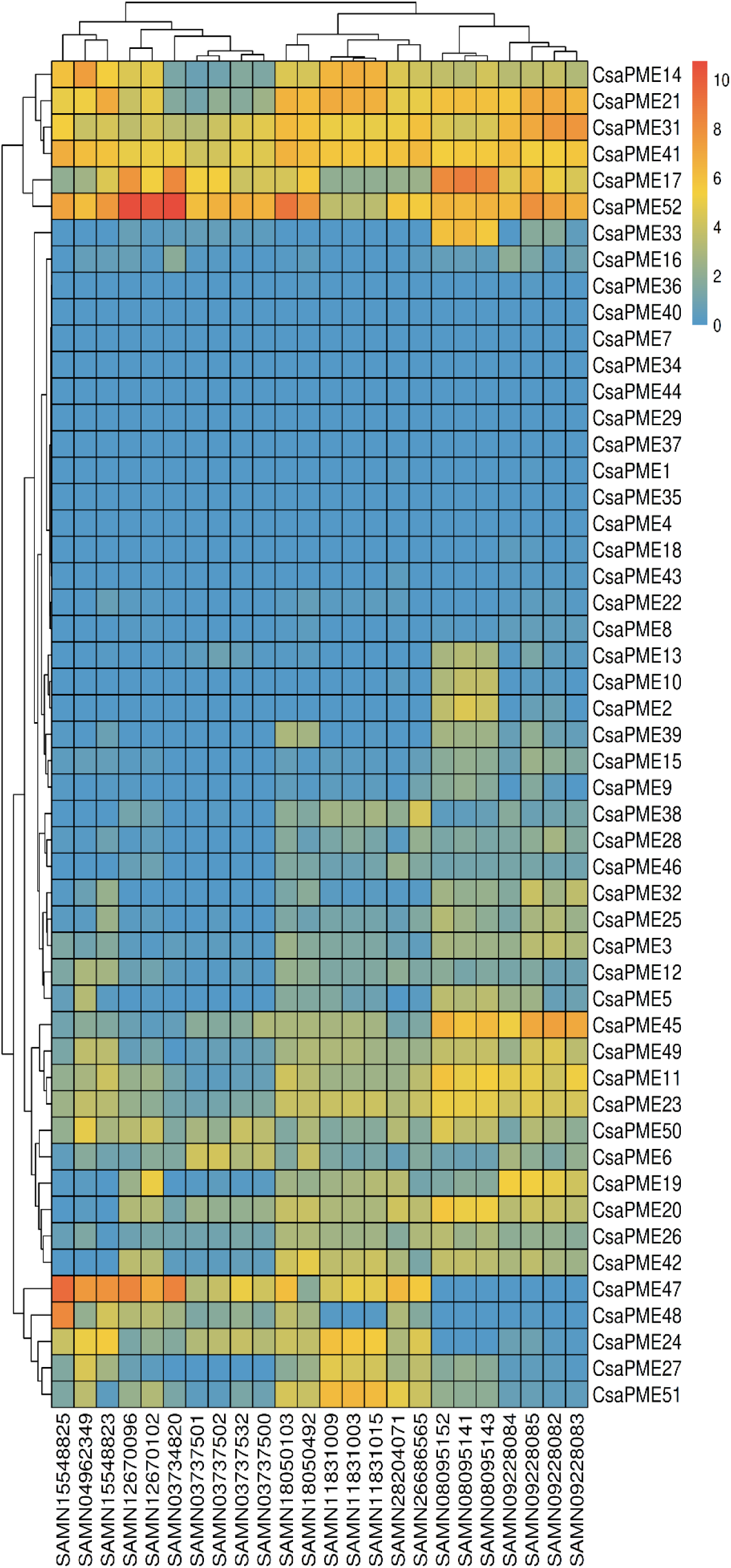
Normalized expression profiles of cucumber PME genes in response to diverse biotic stresses, including fungal, bacterial, viral, and nematode pathogens. Expression data were retrieved from publicly available RNA-seq datasets representing multiple tissues (leaves, roots, and cotyledons) and different post-inoculation time points. The heatmaps were generated using R packages pheatmap, grid, and gridExtra, allowing for a comparative assessment of PME gene regulation during host-pathogen interactions. Please refer to Table S17 for sample identifiers and the corresponding BioSample accessions and experimental conditions. The color intensity represents relative PME gene expression levels across abiotic stress treatments.

**Figure 8:**
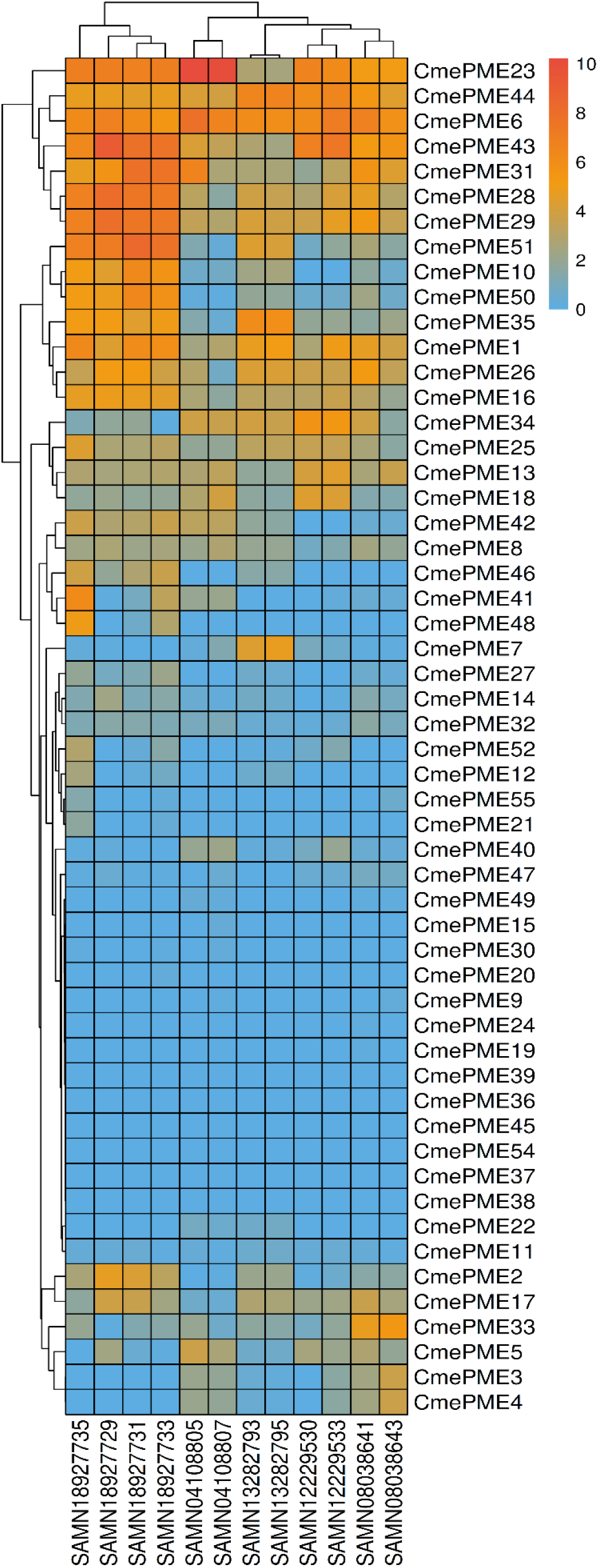
Normalized expression profiles of muskmelon PME genes across multiple abiotic stresses. Expression data were retrieved from publicly available RNA-seq datasets, and heatmaps were generated using R packages pheatmap, grid, and gridExtra. Please refer to Table S18 for sample identifiers and the corresponding BioSample accessions and experimental conditions. The color intensity represents relative PME gene expression levels across abiotic stress treatments.

Several PME genes displayed strong and stress-specific induction. *CsaPME47* and *CsaPME52* emerged as major stress-responsive genes, showing consistent upregulation across diverse abiotic treatments, including H₂S-regulated salt stress, cold stress, drought, heat stress, silicon-mediated salt stress, and waterlogging. For instance, *CsaPME47* and *CsaPME52* were strongly induced in leaves of the plants treated with 200 mM NaCl + 15 µM NaHS, whereas *CsaPME14* and *CsaPME47* were highly upregulated under cold stress in grafted plants. Under drought stress, *CsaPME47* and *CsaPME41* were preferentially induced, while *CsaPME52* and *CsaPME21* showed strong induction under heat and salt-silicon treatments. Tissue-specific responses were also evident, with *CsaPME17* being highly expressed in roots under long-term waterlogging and salt stress, and *CsaPME11*, *CsaPME17*, and *CsaPME52* showing enhanced expression in hypocotyls during waterlogging **(Table S16; Figure 6**). In response to biotic stresses, PME genes exhibited pronounced and dynamic expression patterns. Infection with cucumber green mottle mosaic virus (CGMMV) resulted in strong upregulation of *CsaPME47*, *CsaPME48*, and *CsaPME52* at 20 days post-inoculation, whereas early infection (3 dpi) predominantly induced *CsaPME47* and *CsaPME52*. These genes were highly induced in susceptible lines challenged with *A. cucumerina*. Similarly, during *P. cubensis* infection, *CsaPME52* showed consistently high expression across multiple time points, while other PMEs displayed transient induction. Bacterial infection with *Pseudomonas syringae* pv. *lachrymans* also resulted in strong upregulation of *CsaPME52* **(Table S17; Figure 7**).

Root-specific induction was observed in response to soil-borne pathogens, where *CsaPME17*, *CsaPME52*, and *CsaPME33* were highly expressed during *Meloidogyne incognita* infection, with *CsaPME33* exhibiting nematode-specific induction. Likewise, infection with *F. oxysporum* led to strong upregulation of *CsaPME52*, *CsaPME31*, and *CsaPME45*, with *CsaPME45* being specifically induced under Fusarium and nematode infections. Collectively, these results highlight *CsaPME47* and *CsaPME52* as central hubs in cucumber stress responses, while also revealing pathogen- and tissue-specific roles for selected PME genes.

PME expression profiles were also examined in *C. melo*, and out of 56 PME genes, 24 genes exhibited variable expression patterns across multiple abiotic and biotic stress conditions (**Figures 9-10**). These included 14 Type I PMEs and 10 Type II PMEs, indicating substantial involvement of both PME classes in stress responses. Under abiotic stresses, eight Type I and nine Type II PMEs were highly expressed, whereas 11 Type I and seven Type II PMEs were induced under biotic stress conditions. Notably, five Type I and six Type II PMEs showed overlapping expression under both abiotic and biotic stresses, suggesting shared regulatory mechanisms and multifunctional roles **(Table S18-S19)**. Under waterlogging stress, distinct temporal and tissue-specific expression patterns were observed in hypocotyls. *CmePME43* showed strong early induction at 6 h, followed by progressive downregulation at later time points (24-72 h). A similar early-responsive pattern was observed for *CmePME28*, whereas *CmePME31* and *CmePME51* exhibited delayed induction, with peak expression at 24-48 h and reduced expression at 6 and 72h. Interestingly, *CmePME41* and *CmePME48* were specifically induced during waterlogging and were not responsive to other abiotic stresses. Among these, *CmePME41* also showed strong induction in the roots of susceptible lines following infection with *P. capsici* and *F. oxysporum*, whereas *CmePME48* appeared to be waterlogging-specific and was not induced under biotic stresses.

**Figure 9:**
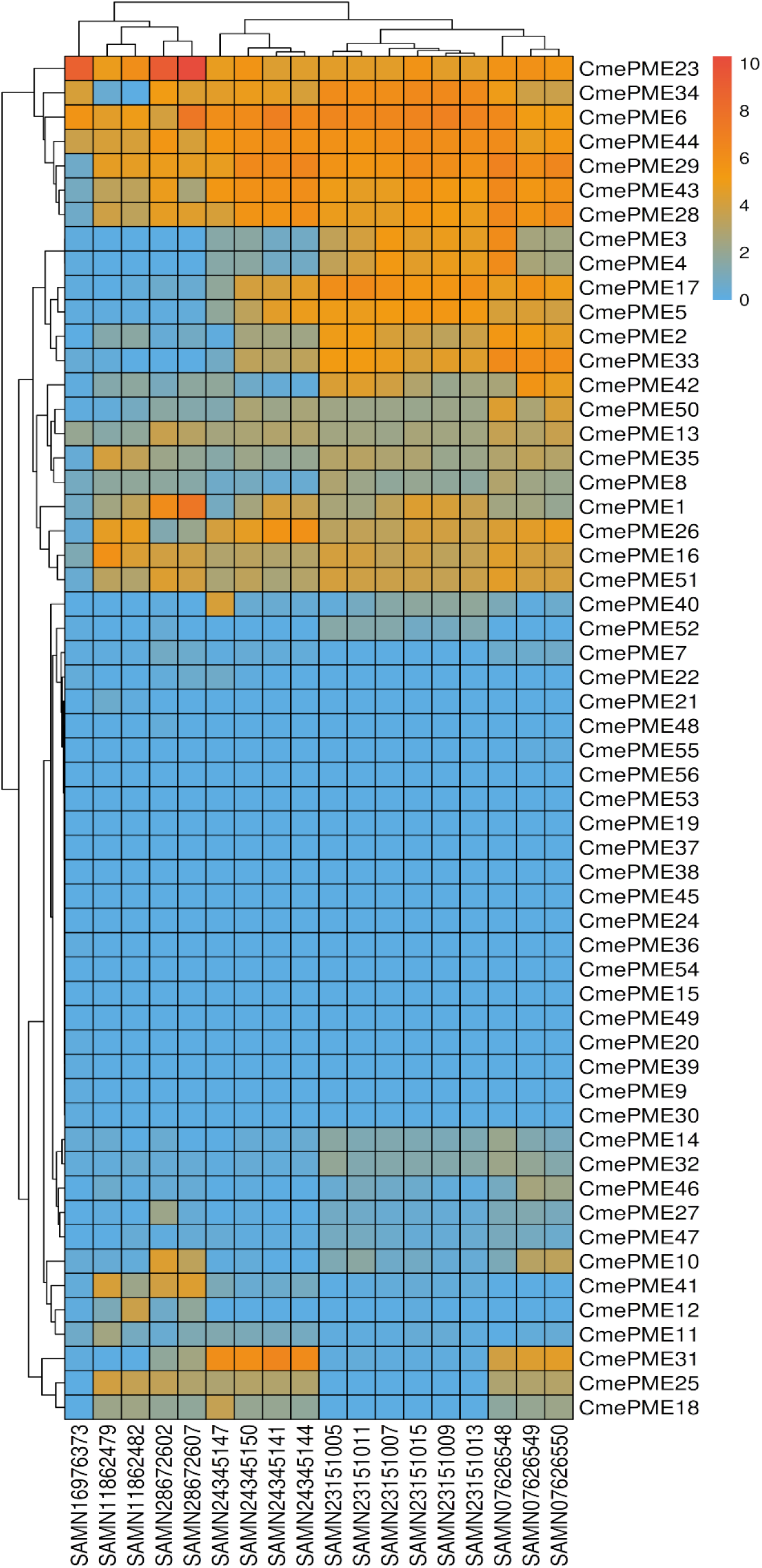
Normalized expression profiles of muskmelon PME genes in response to diverse biotic stresses, including fungal, oomycete, and viral pathogens, as well as mock-inoculated controls. Expression data were retrieved from publicly available RNA-seq datasets representing different post-inoculation time points. The heatmaps were generated using R packages pheatmap, grid, and gridExtra, allowing for a comparative assessment of PME gene regulation during host-pathogen interactions. Please refer to Table S19 for sample identifiers and the corresponding BioSample accessions and experimental conditions. The color intensity represents relative PME gene expression levels across abiotic stress treatments.

Salt stress elicited cultivar-specific responses, with *CmePME23* showing strong induction in leaves of the muskmelon cultivars *Bing XueCui* (BXC) and *Yu Lu* (YL), followed by *CmePME6*. In contrast, *CmePME31* displayed high expression exclusively in BXC, indicating genotype-dependent regulation. Similarly, seed transcriptome analysis under abscisic acid (ABA) and melatonin treatments revealed strong upregulation of *CmePME44*, *CmePME6*, and *CmePME35*, along with exclusive induction of *CmePME7*, which was not expressed under any other abiotic or biotic condition, suggesting a seed-specific regulatory role. Exposure to chilling and cold stress resulted in prominent induction of *CmePME43*, *CmePME6*, *CmePME44*, and *CmePME23* in leaves. Notably, *CmePME34* and *CmePME33* showed stress-specific expression, being induced exclusively under chilling and cold stress, respectively, highlighting specialized roles during low-temperature responses **(Table S18; Figure 8**).

In response to biotic stresses, PME genes exhibited strong pathogen- and tissue-specific expression. Infection with gummy stem blight fungus (*S. cucurbitacearum*) led to high expression of *CmePME23* and *CmePME6* in leaves, while *P. capsici* infection in roots induced *CmePME23*, *CmePME16*, and *CmePME6*. During *F. oxysporum* infection, *CmePME23* showed strong early induction at 12 h and very high expression at 72 h, followed by *CmePME6* and *CmePME1*, indicating sustained activation during disease progression. Likewise, during powdery mildew infection, *CmePME31* exhibited a strong early induction at 1 dpi, followed by gradual downregulation, whereas *CmePME6*, *CmePME29*, *CmePME43*, *CmePME28*, and *CmePME44* were consistently highly expressed across all time points. Infection with tomato leaf curl New Delhi virus (ToLCNDV) resulted in strong induction of *CmePME6*, *CmePME34*, *CmePME44*, and *CmePME29*, whereas *P. xanthii* specifically induced *CmePME31*, in addition to *CmePME29*, *CmePME43*, *CmePME28*, *CmePME6*, and *CmePME44*, highlighting pathogen-specific PME activation **(Table S19; Figure 9**).

To further evaluate whether stress-responsive PME genes also retain structural conservation, representative *Arabidopsis* and rice PMEs, along with their syntenic orthologs in cucumber and muskmelon, were subjected to a three-dimensional structural comparison. Homology models generated using SWISS-MODEL showed high structural similarity between *Arabidopsis* PMEs and their cucurbit counterparts. Structural superposition of *AtPME3* with *CsaPME24*, *CsaPME52*, *CmePME16*, and *CmePME7* revealed very low RMSD values (0.284–0.345 Å), indicating near-identical folding **(Figure S14a)**. Similarly, *AtPME6* and its orthologs *CsaPME38* and *CmePME42* also exhibited low RMSD values (0.449–0.544 Å) upon superposition **(Figure S14b)**. *AtPME18* also showed strong structural congruence with *CsaPME45*, *CmePME34*, and the rice PME LOC_Os01g57854 with RMSD values of 0.476–0.649 Å (**Figure S14c)**. Structural alignment of *AtPME17*-like Type II PMEs and their cucurbit orthologs demonstrated conserved proPME architectures with RMSD values ranging from 0.424 to 0.494 Å **(Figure S14d)**. In contrast, *AtPME34* orthologs displayed comparatively higher structural variation, particularly *CmePME28* (RMSD = 1.433 Å), while other paralogs remained highly similar **(Figure S14e)**.

Collectively, these results demonstrate the extensive temporal, tissue-specific, and pathogen-dependent regulation of PME genes in muskmelon, with *CmePME6* and *CmePME23* emerging as central stress-responsive genes. In contrast, several PMEs (e.g., *CmePME7*, *CmePME34*, *CmePME33*, and *CmePME48*) exhibited highly specialized and condition-specific expression patterns.

## 4. Discussion

Despite the availability of well-annotated genome sequences for cucumber and muskmelon, systematic and comprehensive analyses of their pectin methylesterase gene families remain limited. In this study, we performed a genome-wide identification and characterization of PME genes in both species and investigated their transcriptional responses under biotic and abiotic stress conditions. In the present study, we identified a total of 52 PME genes in cucumber (CsaPMEs) and 56 PME genes in muskmelon (*CmePMEs*). The PME family in both species was dominated by Type I PMEs, which accounted for 58% of the total PME complement in cucumber and 55% in muskmelon. In contrast, Type II PMEs represented 42% and 45%, respectively. This distribution contrasts with previous reports in *A. thaliana* and *O. sativa*, where Type II PMEs constitute a larger proportion of the PME family, accounting for approximately 65% and 52% of the total PMEs, respectively (Jeong et al. 2015; Huang et al. 2022). The relatively higher abundance of Type I PMEs in the cucurbit species may reflect lineage-specific expansion or functional specialization, potentially linked to distinct cell wall remodeling requirements during growth and stress adaptation of these crops.

The uneven chromosomal distribution of PME genes in cucumber and muskmelon can be primarily explained by lineage-specific gene duplication mechanisms acting during cucurbit genome evolution. Our duplication analysis indicates that dispersed duplication was the predominant driver of PME family expansion in both species, accounting for 75% of PME genes in cucumber and 64% in muskmelon. This predominance of dispersed duplication is consistent with the widespread scattering of PME genes across all chromosomes rather than their confinement to a few tandem clusters. Similar observations have been reported in tea, rice, and *Arabidopsis*, where unequal chromosomal distributions of PMEs have been attributed to historical gene and segmental duplication events (Huang et al. 2022). However, unlike *Arabidopsis* and rice, where whole-genome duplication (WGD) and tandem duplication contributed substantially to PME expansion, cucumber and muskmelon show relatively limited contributions from WGD and tandem duplication, suggesting that dispersed duplication played a central role in shaping PME genomic organization in cucurbits. The comparatively higher proportion of proximal duplications observed in muskmelon further indicates species-specific evolutionary trajectories that may underlie functional diversification of PME genes.

Most Type I PMEs contained 2-3 exons, whereas Type II PMEs exhibited greater structural complexity, with 2-4 exons in cucumber and 4-5 exons in muskmelon. The marked variation in intron content among cucumber and muskmelon PME genes indicates substantial structural diversification between the cucumber and muskmelon PME families. Similar to previous reports (Carbone et al. 2025), most PMEs retained conserved catalytic domains, while a subset exhibited expanded gene structures, reflecting directional evolution and functional specialization (Li et al. 2016). The higher intron richness observed in several Type II PMEs may facilitate regulatory flexibility and stress-responsive expression. Conserved motif analysis revealed strong preservation of core PME catalytic motifs across both species, underscoring functional conservation. Notably, the presence of PMEI-associated motifs in Type I PMEs supports the acquisition of regulatory N-terminal domains during evolution. Collectively, these findings suggest that the PMEs of cucumber and muskmelon may have evolved through a balance of structural conservation and gene architecture diversification, supporting adaptive roles under diverse physiological and stress conditions. This conservation preserves the core functional roles of the gene family while promoting functional diversification and reducing the selection pressure (Li et al. 2016).

Phylogenetic analysis revealed a clear evolutionary separation between Type I and Type II PMEs, consistent with previous studies that have shown that PME diversification closely follows domain architecture in *Arabidopsis*, cotton, and tea, among others (Huang et al. 2022). At the same time, the presence of a few exceptions and species-specific expansions highlights the complex and dynamic evolutionary history of the PME gene family across monocots and dicots (Wang et al. 2013; Duan et al. 2016; Li et al. 2016). The subdivision of both PME types into multiple clades reflects lineage-specific diversification and functional specialization. Notably, Clade I-f was characterized by genes exhibiting high intron retention, suggesting recent evolutionary divergence or relaxed structural constraints within this subgroup (Wang et al. 2021). Together, these patterns underscore the interplay between conserved catalytic function and structural innovation in shaping PME evolution.

The abundance and diversity of *cis*-acting regulatory elements in PME promoters suggest that PME gene expression in cucumber and muskmelon is subject to multilayered transcriptional control. The strong enrichment of light- and phytohormone-responsive promoter elements suggests integration of PME activity with developmental cues and hormonal signaling pathways, consistent with the known roles of pectin modification in cell wall remodeling and growth (Wang et al. 2013; Wu et al. 2018). Notably, the predominance of stress-responsive elements, particularly drought-associated MYC and anaerobic-responsive ARE motifs, supports a central role for PMEs in adapting to both abiotic and biotic stresses (Cheng et al. 2022; Lin et al. 2025). The comparatively limited but gene-specific enrichment of growth- and seed-related elements, such as the AAGAA motif in *CmePME20*, further implies functional specialization of individual PME members during development. The strong concordance between promoter architecture and expression profiles further supports the regulatory relevance of stress-associated *cis*-elements in cucumber PMEs. For instance, *CsaPME47* harbors a high density of low-temperature–responsive (LTR) and general stress-responsive regulatory elements, which is consistent with its pronounced induction under cold stress (5 °C) as well as its strong upregulation during drought and heat stress (42 °C) (Wu et al. 2017). This suggests that *CsaPME47* functions as a broad-spectrum stress-responsive PME, likely integrating multiple environmental signals through its complex promoter composition.

Similarly, defense- and stress-related TC-rich repeat elements were associated with pathogen-responsive PME genes. *CsaPME11*, enriched in TC-rich repeats, showed strong induction in roots following infection with *F. oxysporum* and *M. incognita*, while *CsaPME19* and *CsaPME24* were markedly upregulated upon *F. oxysporum* inoculation in roots (Klosterman et al. 2011). Moreover, *CsaPME24* exhibited robust expression in leaves challenged with *P. cubensis*, cucumber green mottle mosaic virus, and powdery mildew fungus, as well as in cotyledons of susceptible lines infected with *P. xanthii* (Falade 2021; Meng et al. 2022). Collectively, these observations suggest that the enrichment of defense-related *cis*-elements, particularly TC-rich repeats, underpins the pathogen- and tissue-specific activation of PME genes, highlighting their crucial roles in cucumber stress and immune responses. Moreover, *CsaPME5* and *CsaPME45* contained the highest number of stress-responsive *cis*-regulatory elements, indicating strong transcriptional responsiveness to environmental cues (Cheng et al. 2022; Lin et al. 2025). Consistent with this promoter enrichment, both genes showed marked upregulation under waterlogging stress in hypocotyl tissues, suggesting a role in hypoxia-associated cell wall remodeling (Arora et al. 2017; Zhu et al. 2023). Notably, *CsaPME45* displayed a broader stress-responsive expression profile, being strongly induced not only in hypocotyls but also in roots under prolonged waterlogging, as well as in roots exposed to salt stress for three days. In addition to abiotic stress responses, *CsaPME45* was also highly expressed under biotic stress, particularly in roots infected with *M. incognita* and *F. oxysporum* (Klosterman et al. 2011).

In the case of muskmelon, several PME genes also exhibited a strong correspondence between stress-responsive *cis*-element enrichment and transcriptomic activation under stress conditions. *CmePME2*, which harbors a high density of stress-responsive regulatory elements, was markedly upregulated in the hypocotyl under waterlogging stress (Arora et al. 2017), as well as in leaves of susceptible plants following powdery mildew infection and ToLCNDV inoculation, indicating its involvement in both abiotic and biotic stress responses. Similarly, *CmePME3* and *CmePME42* possessed abundant stress-responsive *cis*-elements and were consistently upregulated during ToLCNDV infection and powdery mildew challenge (Risk et al. 2013), suggesting shared regulatory control during pathogen attack. Notably, *CmePME17* contained the highest abundance of stress-associated elements, including STRE, ARE, MYC, and TC-rich repeats, and showed strong induction specifically under biotic stresses such as powdery mildew and ToLCNDV, but not under abiotic treatments, implying a specialized role in pathogen-mediated defense. In contrast, *CmePME10*, which also carries multiple stress-responsive elements, exhibited pronounced upregulation under waterlogging stress and *F. oxysporum* infection (Sella et al. 2016; Zhang et al. 2021), highlighting the functional diversification of muskmelon PMEs in coordinating responses to distinct environmental and pathogenic cues.

The PPI network analysis showed that PMEs in both cucumber and muskmelon could serve as integral components of a highly coordinated cell wall-modifying machinery, rather than acting as isolated enzymes. The dense interaction networks, enriched for pectate lyases, polygalacturonases, glycosyl hydrolases, and PME inhibitors, support the concept that pectin methylesterification and depolymerization are tightly coupled processes, ensuring precise regulation of cell wall remodeling during growth and stress adaptation. The enrichment of extracellular localization further corroborates the central role of PMEs in apoplastic cell wall dynamics. Importantly, integration of PPI topology with transcriptomic data revealed that several hub- and top-interacting PMEs are also transcriptionally highly responsive to stress, reinforcing their functional relevance. In cucumber, *CsaPME41*, a highly connected PPI node, showed strong and consistent upregulation across multiple abiotic (drought, heat, salt, silicon-mediated salt stress, and waterlogging) as well as biotic stresses (Sella et al. 2016; Zhang et al. 2021), suggesting that this PME may act as a core regulator linking cell wall remodeling to broad stress responses. Similarly, other highly interacting PME-associated proteins in cucumber were transcriptionally induced in response to abiotic stresses, such as drought and heat (Wu et al. 2017; Cheng et al. 2022), as well as biotic challenges including *P. cubensis* and viral infections, indicating the coordinated activation of the PME-centered network under diverse stress conditions.

A comparable pattern was also observed in muskmelon, where *CmePME23*, identified among the highly connected PME-associated proteins, emerged as a major stress-responsive gene, showing strong induction during biotic stresses, particularly *F. oxysporum* infection, and across abiotic stresses, with maximal expression under salt treatment (Yan et al. 2018; Liu et al. 2018). Additional top PPI-associated PMEs in muskmelon were also transcriptionally activated during powdery mildew infection, *P. capsici* infection, and *F. oxysporum* infection, as well as waterlogging, chilling, and NaCl treatments (Yan et al. 2018), highlighting the conserved yet species-specific deployment of PME interaction networks in stress adaptation. These findings suggest that hub PMEs identified through PPI analysis represent key regulatory nodes that integrate cell wall remodeling with environmental and pathogen-derived signals.

The syntenic conservation of PME genes between cucumber and muskmelon reflects their close phylogenetic relationship. It indicates that the PME gene repertoire has remained largely stable following speciation within the *Cucumis* lineage. Such conservation suggests that many PME loci are under functional constraint, likely due to their essential roles in cell wall remodeling and development (Wu et al. 2018). The moderate collinearity observed with *A. thaliana* implies that a core set of PME genes predates the divergence of major dicot lineages and has been selectively retained, supporting ancient and conserved PME functions across dicots. In contrast, the limited synteny with rice is consistent with the monocot-dicot split and highlights extensive lineage-specific rearrangements and diversification of PME genes in monocots. Evidence for functional conservation in cucurbits is further supported by synteny with *AtPME58* (AT5G49180), a key regulator of homogalacturonan modification during seed coat mucilage extrusion (Turbant et al. 2016). In cucumber, two duplicated *AtPME58*-like orthologs (*CsaPME18* and *CsaPME42*) were retained, while muskmelon harbors corresponding duplicated counterparts (*CmePME24* and *CmePME52*). Notably, all five genes cluster tightly within the same phylogenetic clade, indicating a shared evolutionary origin and conserved domain architecture. The retention of multiple syntenic copies of PME genes suggests potential subfunctionalization or dosage-sensitive regulation of pectin remodeling processes in cucurbits. However, experimental validation will be necessary to resolve the functional divergence among these paralogs (Wang et al. 2021; Zhang et al. 2022, 2024).

The ratio of nonsynonymous to synonymous substitutions (Ka/Ks) is a robust indicator of the selective forces shaping gene family evolution. Ka/Ks analysis of 49 one-to-one orthologous PME gene pairs between the cucurbit species revealed that all PME orthologs in both cucumber and muskmelon exhibited Ka/Ks values well below 1. This pattern provides strong evidence that PME genes have evolved predominantly under purifying selection, indicating strong evolutionary constraints acting to preserve their protein sequences and core biological functions. Similar evolutionary trends have been reported across diverse plant lineages. Genome-wide studies in the tea plant (Huang et al. 2022), cotton (Li et al. 2016), persimmon (Zhang et al. 2022), pear (Zhang et al. 2024), and walnut (Qin et al. 2024) have consistently shown Ka/Ks ratios < 1 for the majority of PME gene pairs, reflecting stabilizing selection during evolution. Although a few lineage-specific PME genes in pear showed signatures of positive selection, such cases appear to be rare and exceptional. The comparative expression analysis revealed both conservation and diversification of PME-mediated stress responses in cucumber and muskmelon. While stress-responsive PME activation is conserved in both species, cucumber responses were primarily dominated by Type I PMEs. In contrast, muskmelon showed a more balanced involvement of Type I and Type II PMEs, indicating evolutionary divergence in regulatory strategies. The consistent induction of *CsaPME47*/*CsaPME52* and *CmePME6*/*CmePME23* across multiple stresses suggests that these genes function as core hubs in cell wall remodeling during stress adaptation. Tissue- and pathogen-specific induction patterns further support functional specialization of individual PMEs in root-pathogen interactions and disease progression. Moreover, distinct early and late induction dynamics highlight the role of PMEs in both stress perception and downstream acclimation processes.

The functional relevance of stress-responsive cucurbit PMEs is further supported by their evolutionary conservation with experimentally characterized *Arabidopsis* homologs. A notable example is *AtPME3* (AT3G14310), a ubiquitously expressed pectin methylesterase with well-established roles in cell wall remodeling, root development, and defense-related processes (Guénin et al. 2017; Song 2022). *AtPME3* has been shown to modulate pectin methylesterification at host-parasite interfaces, thereby influencing susceptibility to the parasitic plant *Phelipanche ramosa*, as well as acting as a susceptibility factor during infections by *Pectobacterium carotovorum* and *Botrytis cinerea*. Loss-of-function studies demonstrated that altered PME activity and reduced pectin methylesterification significantly affect pathogen or parasite establishment, highlighting *AtPME3* as a key regulator of cell wall-mediated stress responses (Grandjean et al. 2024). In the present study, *CsaPME24* in cucumber and CmPME16 in muskmelon were identified as syntenic orthologs of *AtPME3* and clustered within the same phylogenetic clade, suggesting conservation of ancestral function. Consistent with the known stress-related roles of *AtPME3*, both cucurbit orthologs displayed broad and robust induction under multiple biotic and abiotic stresses. *CsaPME24* was strongly upregulated in response to diverse abiotic stresses, including salt (Yan et al. 2018) and silicon treatment, heat (Huang et al. 2017), and drought, as well as during infections by *P. xanthii* and *P. cubensis* in susceptible tissues (Meng et al. 2022). Similarly, *CmPME16* showed pronounced induction during waterlogging stress in hypocotyls and under multiple biotic challenges, including powdery mildew, tomato leaf curl New Delhi virus (ToLCNDV), and *P. capsici*, particularly in susceptible genotypes (Feng et al. 2010).

*AtPME6*/*HMS* (AT1G23200), a pectin methylesterase implicated in embryo development, seed mucilage modification, and stomatal movement, has also been shown to exhibit consistent upregulation during parasitic haustorium formation in *A. thaliana*, indicating a potential role in pectin remodeling and host defense responses during plant–parasite interactions (Levesque-Tremblay et al. 2015; Leso et al. 2023). In cucumber and muskmelon, the syntenic orthologs *CsaPME38* and CmPME42, which clustered closely with *AtPME6* in phylogenetic analysis, exhibited highly specific induction exclusively during powdery mildew (*P. xanthii*) infection in susceptible tissues (Meng et al. 2022). This restricted pathogen-specific expression suggests that these PMEs exhibit functional specialization in host-pathogen interface remodeling rather than broad stress responses. The conserved yet narrowly tuned activation pattern implies that *AtPME6*-like PMEs may regulate precise pectin demethylesterification events critical for fungal colonization or defense, warranting targeted functional validation in cucurbit-powdery mildew pathosystems. Another *AtPME18* (AT1G11580), a multifunctional PME implicated in root growth, plant-pathogen interactions, and defense responses, was reported to exhibit ribosome-inactivating protein (RIP) activity following post-translational processing (De-la-Peña et al. 2008; Stefanowicz et al. 2021; Coculo and Lionetti 2022). In cucumber and muskmelon, this gene was represented by syntenic orthologs *CsaPME45* and *CmePME34*, respectively, which also had synteny with a rice PME (LOC_Os01g57854), indicating deep evolutionary retention across monocots and dicots. Consistent with the *Arabidopsis* phenotype, *CsaPME45* showed strong induction in roots under waterlogging and salt stress, and was highly upregulated during biotic stress caused by *F. oxysporum* and *M. incognita* (Klosterman et al. 2011; Goode et al. 2025). Similarly, *CmePME34* was activated during chilling stress and displayed pronounced induction under multiple biotic challenges, including *P. xanthii*, *F. oxysporum*, and ToLCNDV infections.

Additional evidence for functional conservation of defense-associated PMEs was provided by synteny with *AtPME17* (AT2G45220), a Type II PME known to be proteolytically activated by the subtilase SBT3.5 and to play a critical role in jasmonate ethylene-mediated resistance against *Botrytis cinerea* (Sénéchal et al. 2014; Del Corpo et al. 2020). In cucumber and muskmelon, two conserved orthologous pairs (*CsaPME20*/*CsaPME16* and *CmePME22*/*CmePME27*) were identified, which clustered within a single phylogenetic clade, supporting a shared evolutionary origin and a conserved proPME architecture. Notably, among these, *CsaPME20* showed strong and specific induction during *M. incognita* infection in roots, paralleling the pathogen-induced activation of *AtPME17* in *Arabidopsis* (Goode et al. 2025). This selective biotic responsiveness suggests that *AtPME17*-like PMEs may retain specialized roles in root-pathogen interactions in cucurbits, potentially involving regulated PRO-domain processing and localized pectin demethylesterification.

*AtPME34* (AT3G49220) is a guard cell-expressed PME that regulates stomatal movement, transpiration, and thermotolerance through pectin remodeling during heat stress (Huang et al. 2017; Wu et al. 2017; Coculo and Lionetti 2022). In cucumber and muskmelon, two retained orthologous pairs (*CsaPME21*/*CsaPME31* and *CmePME28*/*CmePME45*) were identified, suggesting preservation of *AtPME34*-like functions following cucurbit diversification. Notably, *CsaPME21* displayed broad induction across abiotic stresses, including high-temperature treatment (42 °C), consistent with the heat-induced expression and ABA responsiveness of *AtPME34*, whereas *CsaPME31* showed preferential activation under biotic stresses (Raiola et al. 2011; Liu et al. 2018), indicating subfunctionalization among paralogs. In muskmelon, *CmePME28* exhibited stress-specific induction in hypocotyls under waterlogging, while *CmePME45* did not respond predominantly to pathogen challenges (Zhang et al. 2021), further supporting divergence in regulatory deployment. These patterns suggest that *AtPME34*-associated roles in stomatal and stress-mediated cell wall flexibility are evolutionarily conserved in cucurbits but have diversified into distinct abiotic- and biotic-responsive modules that necessitate experimental validation.

Another PME, *AtPME3* (AT3G14310), is a multifunctional, ubiquitously expressed PME implicated in seed germination, root development, pavement cell morphogenesis, pathogen susceptibility, and metal tolerance through the dynamic regulation of cell wall pectin demethylesterification (Raiola et al. 2011; Guénin et al. 2017). Functional studies have demonstrated that *AtPME3* acts as a susceptibility factor during nematode, bacterial, and necrotrophic fungal infections, and modulates abiotic stress responses, including Zn²⁺ toxicity and Ca²⁺-dependent cell wall cation binding (Weber et al. 2013). In cucumber, two syntenic orthologs (*CsaPME24* and *CsaPME52*) showed marked regulatory divergence, with *CsaPME24* displaying stress-specific induction under drought, heat, salinity, silicon, and selected biotic challenges, whereas *CsaPME52* exhibited broad responsiveness across most abiotic and biotic stresses (Weber et al. 2013; Guénin et al. 2017; Coculo and Lionetti 2022), suggesting partial functional redundancy with regulatory specialization. In muskmelon, *CmePME7* and *CmePME16* further illustrate this divergence, where *CmePME7* responded selectively to ABA and melatonin signaling, while *CmePME16* was predominantly activated under waterlogging and multiple pathogen infections (Gao et al. 2020).

To further substantiate the functional conservation inferred from synteny and stress-responsive expression, we compared the three-dimensional structures of representative *Arabidopsis* and rice PMEs, as well as their cucurbit orthologs, using homology modeling with SWISS-MODEL (https://swissmodel.expasy.org/) (Waterhouse et al. 2018) and structural superposition using Pymol (Schrodinger 2023). Structural superimposition of cucurbit and rice PMEs against *AtPME3* revealed near-identical folding and strong conservation of the catalytic core despite regulatory divergence. Similarly, *AtPME6* and its cucurbit orthologs (*CsaPME38* and *CmePME42*) exhibited the preservation of precise structural features likely required for pathogen-interface pectin remodeling. *AtPME18* and its orthologs from cucumber, muskmelon, and rice also exhibited high structural congruence, consistent with deep evolutionary retention across monocots and dicots. Structural superposition of *AtPME17*-like Type II PMEs revealed conserved proPME architectures, reinforcing their specialized roles in regulated defense responses (Sénéchal et al. 2014; Del Corpo et al. 2020). In contrast, *AtPME34* orthologs exhibited slightly higher structural divergence, particularly in *CmePME28*, consistent with the observed regulatory subfunctionalization among paralogs.

Altogether, the consistently low RMSD values across *Arabidopsis*, cucumber, muskmelon, and rice PMEs indicate strong conservation of three-dimensional structure despite pronounced transcriptional and regulatory divergence. This structural conservation highlights stringent evolutionary constraints on PME catalytic function, while permitting regulatory and functional specialization following gene duplication.

Despite providing a comprehensive evolutionary and stress-responsive framework for PME genes in cucumber and muskmelon, a few unanswered questions remain. For example, which PMEs act as primary regulators versus downstream responders during stress adaptation? How do PMEs coordinate with other cell wall-modifying enzymes to balance growth and defense? Can the selective manipulation of specific PME isoforms enhance stress tolerance without compromising growth? Addressing these questions will be essential for fully harnessing PMEs in cucurbit crop improvement. This study also has a few limitations that need to be acknowledged. First, the analyses were based on *in silico* predictions and meta-transcriptomic datasets derived from heterogeneous experiments, genotypes, tissues, stress types, and intensities, which may not accurately reflect context-specific gene expression and limit direct cross-condition comparisons. Second, protein-protein interaction networks were inferred from orthology-based predictions and require experimental validation. Third, any functional redundancy among the PME genes would complicate the assignment of precise biological roles to individual genes based solely on expression patterns. Future research may therefore prioritize targeted functional validation of key stress-responsive PME genes using genetic approaches such as CRISPR/Cas-mediated knockouts, overexpression, and tissue-specific expression, coupled with biochemical assays of PME activity. Moreover, exploring the natural allelic variation of PME genes across cucurbit germplasm and linking it to stress-resilience traits could accelerate translational breeding applications. Nevertheless, the study reveals stress-responsive cucurbit PMEs as conserved yet adaptable regulators of cell wall plasticity, and positions them as promising molecular targets for enhancing multi-stress resilience in cucurbit crops.

## 5. Conclusions

This study elucidated the evolutionary organization, regulatory landscape, and stress-associated roles of the PME gene family in cucumber and muskmelon. Cucurbit PMEs exhibited a higher proportion of Type I members than model dicots and monocots, reflecting lineage-specific expansion and potential specialization linked to cell wall remodeling. PME family diversification in cucurbits seems to be driven predominantly by dispersed duplication, resulting in species-specific evolutionary patterns while maintaining strong conservation of catalytic domains and motifs. Phylogenetic segregation of Type I and Type II PMEs, coupled with clade-specific intron variation, highlights how conserved enzymatic function coexists with flexible gene architecture, enabling regulatory diversification. Synteny and Ka/Ks analyses further indicate that most PME genes are evolutionarily constrained and maintained under strong purifying selection since the early divergence of dicots. Integrated promoter, protein-protein interaction, and meta-transcriptomic analyses reveal that PMEs operate within coordinated cell wall-modifying networks and are tightly regulated by stress-responsive signals. The concordance between *cis*-regulatory enrichment and stress-induced expression supports a central role of PMEs in cell wall plasticity during abiotic and biotic challenges. Differential deployment of PME Type I dominance in cucumber and balanced Type I/Type II involvement in muskmelon highlights species-specific regulatory strategies. The identification of conserved stress-responsive hub PMEs alongside highly context-specific members suggests functional partitioning within the family. Conserved orthology with functionally characterized *Arabidopsis* PMEs supports the evolutionary retention of stress-related functions of cucurbit PMES, alongside regulatory divergence following duplication. Targeted experimental validation, including functional genomics and biochemical assays, will be essential to confirm the roles of these candidate PMEs and to translate these findings into strategies for enhancing multi-stress resilience in cucurbit crops.

## Acknowledgements

**AKS** acknowledges AcSIR, India, for the Integrated Dual-Degree Program (IDDP) and the research fellowship for IDDP under the CSIR-GATE-JRF program (Award No.: 31/ GATE/11(55)/2025-EMR-I). **BRD** acknowledges the University Grants Commission (UGC), India, for the junior and senior research fellowships, and the Academy of Scientific and Innovative Research (AcSIR), India, for the Ph.D. program enrollment. **SSN** acknowledges AcSIR for enrollment in the Ph.D. program and CSIR-JRF for the research fellowship.

## Author Contributions

**Anand Kumar Shukla**: Conceptualization, Data curation, Formal analysis, Investigation, Methodology, Software, Visualization, Writing-original draft. **Bhakti R. Dayama**: Data curation, Methodology, Supervision, Visualization, Writing-original draft. **Shubham S. Nikam**: Data curation, Investigation, Writing-review and editing. **Narendra Y. Kadoo**: Conceptualization, Funding acquisition, Project administration, Resources, Supervision, Writing-review and editing.

## Competing Interests

The authors have no competing interests to declare that are relevant to the content of this article.

## Funding

This work was supported by the Council of Scientific and Industrial Research (CSIR), New Delhi, India [grant numbers MLP101226 and MLP101326].

## Data availability

The data supporting the findings of this study are presented within the manuscript.

## Generative AI statement

The authors used ChatGPT to enhance the readability of some text. The authors reviewed and edited the content and take full responsibility for the accuracy and integrity of the publication’s content.

## Supplementary Figure Legends

**Figure S1:** Gene structures (exon–intron organization) of 30 cucumber Type I PME genes based on genomic and CDS sequences, with exons represented as boxes and introns as connecting lines. The 3’ and 5’ UTRs are also shown. Type I PME genes are characterized by the presence of both PME and PMEI domains.

**Figure S2:** Gene structures (exon–intron organization) of 22 cucumber Type II PME genes based on genomic and CDS sequences, with exons represented as boxes and introns as connecting lines. The 3’ and 5’ UTRs are also shown. Unlike Type I members, Type II PME genes lack the PMEI domain and contain only the conserved PME catalytic region.

**Figure S3:** Gene structures (exon–intron organization) of 31 muskmelon Type I PME genes based on genomic and CDS sequences, with exons represented as boxes and introns as connecting lines. The 3’ and 5’ UTRs are also shown. Type I PME genes are characterized by the presence of both PME and PMEI domains.

**Figure S4:** Gene structures (exon–intron organization) of 25 muskmelon Type II PME genes based on genomic and CDS sequences, with exons represented as boxes and introns as connecting lines. The 3’ and 5’ UTRs are also shown. Unlike Type I members, Type II PME genes lack the PMEI domain and contain only the conserved PME catalytic region.

**Figure S5:** Schematic representation of the conserved domain organization of cucumber PME proteins. The domains were identified using conserved domain databases, illustrating the presence of the PME catalytic domain and, in some members, the pectin methylesterase inhibitor (PMEI) domain. The relative lengths and positions of domains are shown for each protein, demonstrating their structural conservation and divergence in cucumber.

**Figure S6:** Schematic representation of the conserved domain organization of muskmelon PME proteins. The domains were identified using conserved domain databases, illustrating the presence of the PME catalytic domain and, in some members, the pectin methylesterase inhibitor (PMEI) domain. The relative lengths and positions of domains are shown for each protein, demonstrating their structural conservation and divergence in muskmelon.

**Figure S7:** Chromosomal localization of cucumber PME genes. PME genes were mapped onto individual chromosomes based on their genomic coordinates obtained from the reference genome annotation, illustrating their genomic organization and distribution patterns.

**Figure S8:** Chromosomal localization of muskmelon PME genes. PME genes were mapped onto individual chromosomes based on their genomic coordinates obtained from the reference genome annotation, illustrating their genomic organization and distribution patterns.

**Figure S9:** Conserved motif distribution of cucumber PME proteins identified using MEME analysis. The top ten conserved motifs were mapped onto each protein, revealing conserved motif patterns across the gene family.

**Figure S10:** Conserved motif distribution of muskmelon PME proteins identified using MEME analysis. The top ten conserved motifs were mapped onto each protein, revealing conserved motif patterns across the gene family.

**Figure S11:** STRING-based protein–protein interaction (PPI) networks of PME proteins in (A) cucumber and (B) muskmelon. The networks were constructed to predict functional associations among the PME proteins based on known and predicted interactions. Nodes represent PME proteins, and edges indicate interaction confidence, providing insights into coordinated functional roles of PMEs.

**Figure S12:** Gene Ontology enrichment analysis of cucumber PME interaction network

**Figure S13:** Gene Ontology enrichment analysis of muskmelon PME interaction network

**Figure S14:** Structural superimposition of PME proteins from *Arabidopsis*, cucumber, muskmelon, and rice.

a) Structural superimposition of PME proteins aligned to *AtPME3* (reference). *AtPME3* (red, reference); *CmePME16* (blue, RMSD = 0.284 Å); *CmePME7* (green, RMSD = 0.344 Å); *CsaPME24* (magenta, RMSD = 0.345 Å); *CsaPME52* (cyan, RMSD = 0.345 Å).

b) Structural superimposition of PME proteins aligned to *AtPME6* (reference). *AtPME6* (red, reference); *CsaPME38* (cyan, RMSD = 0.544 Å); *CmePME42* (magenta, RMSD = 0.449 Å).

c) Structural superimposition of PME proteins aligned to *AtPME18* (reference). *AtPME18* (red, reference); *CmePME34* (green, RMSD = 0.476 Å); *CsaPME45* (magenta, RMSD = 0.478 Å); Loc_Os01g57854 (orange, RMSD = 0.649 Å).

d) Structural superimposition of PME proteins aligned to *AtPME17-*like (reference). *AtPME17*-like (red, reference); *CsaPME16* (magenta, RMSD = 0.494 Å); *CsaPME20* (cyan, RMSD = 0.483 Å); *CmePME22* (blue, RMSD = 0.424 Å); *CmePME27* (green, RMSD = 0.488 Å).

e) Structural superimposition of PME proteins aligned to *AtPME34* (reference). *AtPME34* (red); *CmePME28* (green, RMSD = 1.433 Å); *CmePME45* (blue, RMSD = 0.460 Å); *CsaPME21* (magenta, RMSD = 0.483 Å); *CsaPME31* (cyan, RMSD = 0.459 Å).

## Supplementary Table Legends

**Table S1**: Physicochemical properties of pectin methylesterase (PME) family proteins identified in cucumber (*Cucumis sativus*)

**Table S2**: Physicochemical properties of pectin methylesterase (PME) family proteins identified in muskmelon (*Cucumis melo*)

**Table S3**: Intron–exon organization of PME family genes identified in cucumber (*Cucumis sativus*). Intron length was calculated as gene length − CDS length, and intron percentage was computed as (intron length / gene length) × 100

**Table S4**: Intron–exon organization of PME family genes identified in muskmelon (*Cucumis melo*). Intron length was calculated as gene length − CDS length, and intron percentage was computed as (intron length / gene length) × 100

**Table S5**: Simple Enrichment Analysis (SEA) of conserved motifs identified in cucumber PME proteins, highlighting statistically enriched motifs and their associated functional significance

**Table S6**: Simple Enrichment Analysis (SEA) of conserved motifs identified in muskmelon PME proteins, highlighting statistically enriched motifs and their associated functional significance

**Table S7**: InterProScan-based functional annotation of conserved motifs identified in cucumber PME proteins

**Table S8**: InterProScan-based functional annotation of conserved motifs identified in Muskmelon PME proteins

**Table S9**: Distribution of cis-acting regulatory elements in the promoter regions of cucumber PME genes. The identified elements were classified into four major functional categories: (i) light-responsive, (ii) phytohormone-responsive, (iii) stress-responsive, and (iii) growth and development-related elements

**Table S10**: Distribution of cis-acting regulatory elements in the promoter regions of muskmelon PME genes. The identified elements were classified into four major functional categories: (i) light-responsive, (ii) phytohormone-responsive, (iii) stress-responsive, and (iii) growth and development-related elements

**Table S11**: CytoHubba-based network analysis of cucumber PME proteins to identify hub genes based on topological parameters within the protein–protein interaction network

**Table S12**: CytoHubba-based network analysis of muskmelon PME proteins to identify hub genes based on topological parameters within the protein–protein interaction network

**Table S13**: Gene duplication analysis of PME family genes across four plant species: *Arabidopsis thaliana*, *Oryza sativa*, *Cucumis sativus*, and *Cucumis melo*, highlighting duplication modes and evolutionary relationships

**Table S14**: PME gene IDs and their corresponding duplication types identified across *Arabidopsis thaliana*, *Oryza sativa*, *Cucumis sativus*, and *Cucumis melo*

**Table S15**: Synteny analysis of PME family genes among cucumber, muskmelon, *Arabidopsis thaliana*, and rice, illustrating conserved genomic blocks and orthologous gene relationships

**Table S16**: Normalized expression profiles of cucumber PME genes across various abiotic stress conditions based on transcriptomic data

**Table S17**: Normalized expression profiles of cucumber PME genes across various biotic stress conditions based on transcriptomic data

**Table S18**: Normalized expression profiles of muskmelon PME genes across various abiotic stress conditions based on transcriptomic data

**Table S19**: Normalized expression profiles of cucumber PME genes across various biotic stress conditions based on transcriptomic data

## Notes

### Competing Interest Statement

The authors have declared no competing interest.

