## Supplementary figures and images for "Genome-wide, evolutionary, and stress-responsive landscape of the Pectin methylesterase gene family in cucumber and muskmelon"

### Figure S1

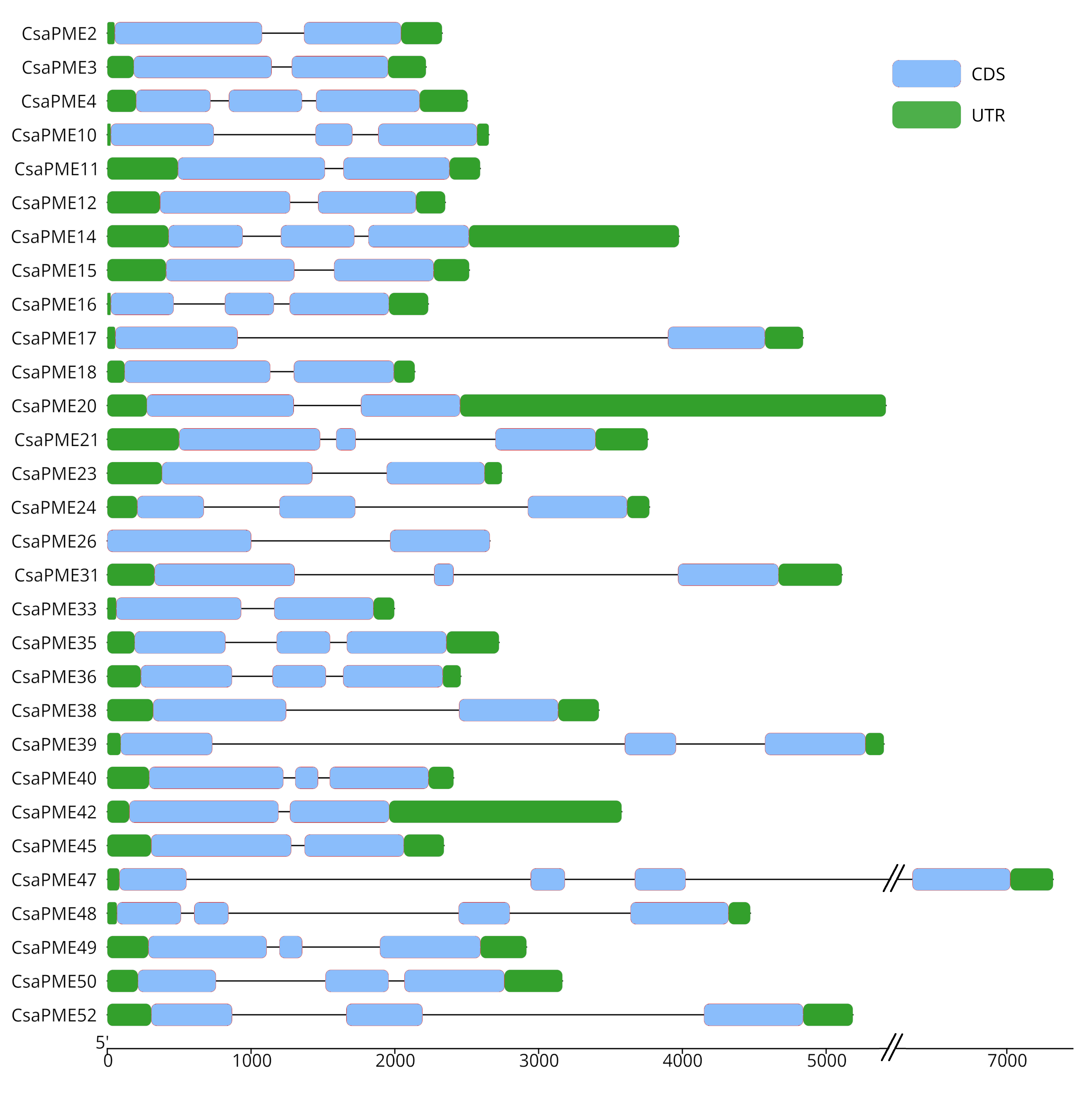

### Figure S2

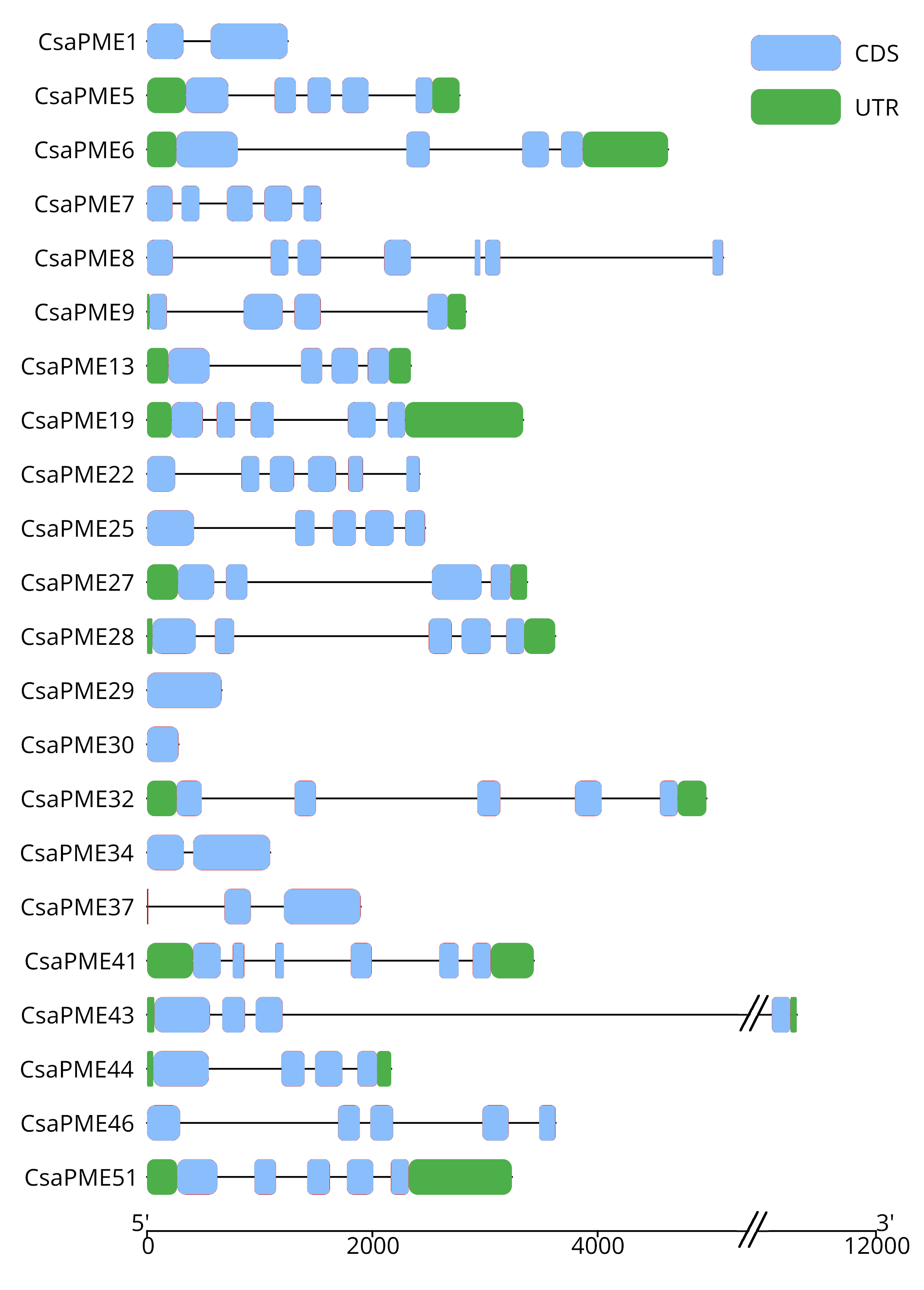

### Figure S3

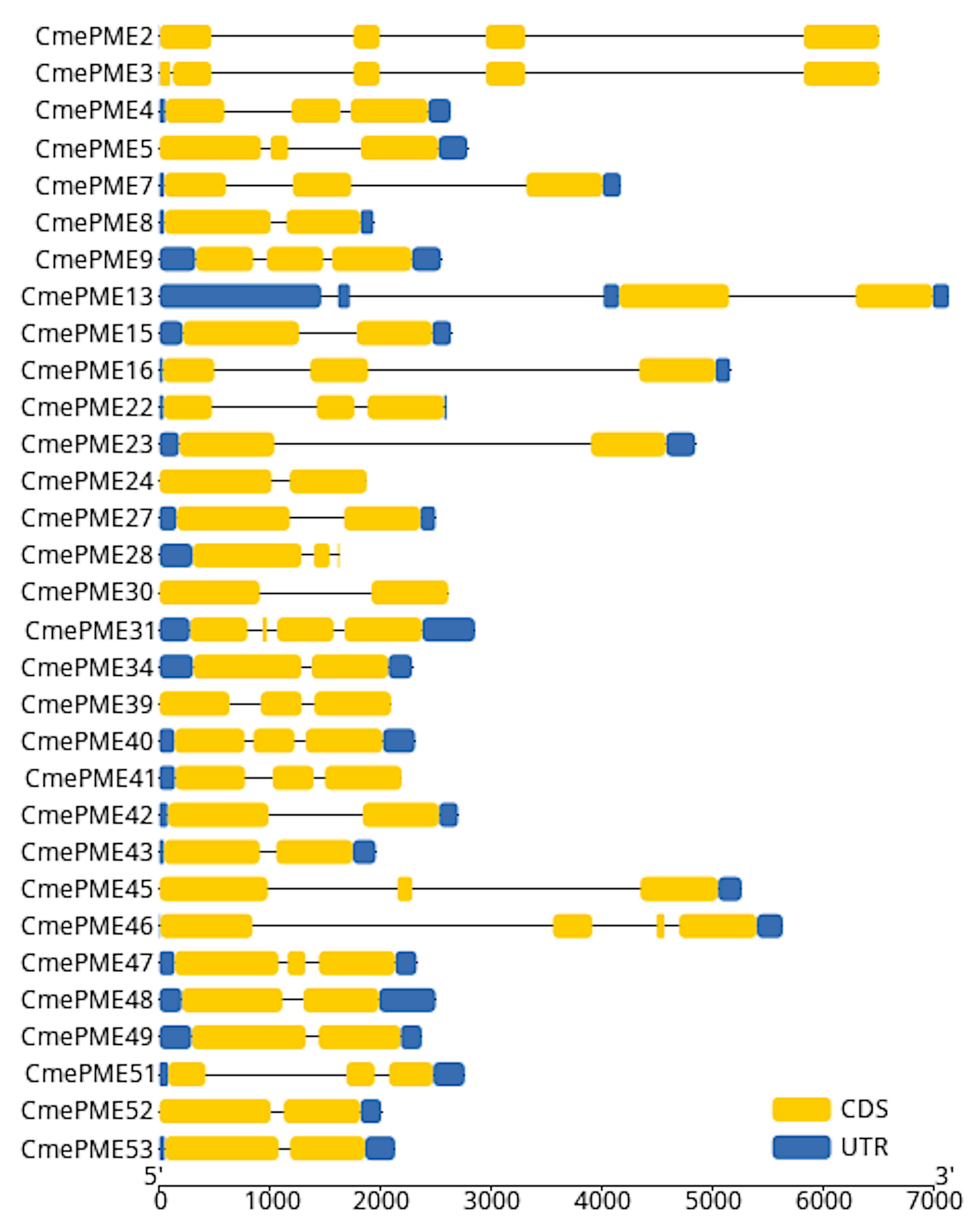

### Figure S4

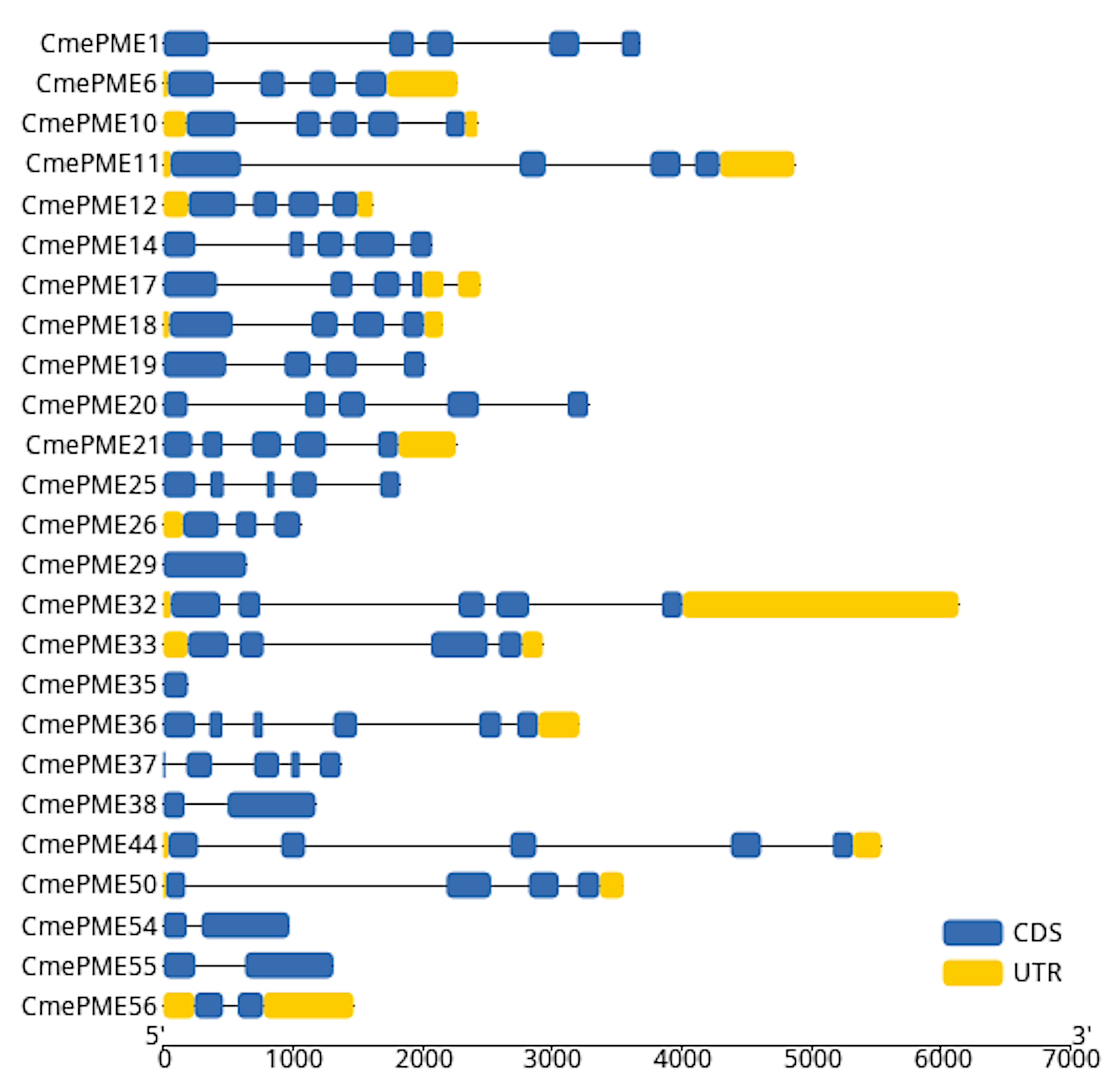

### Figure S5

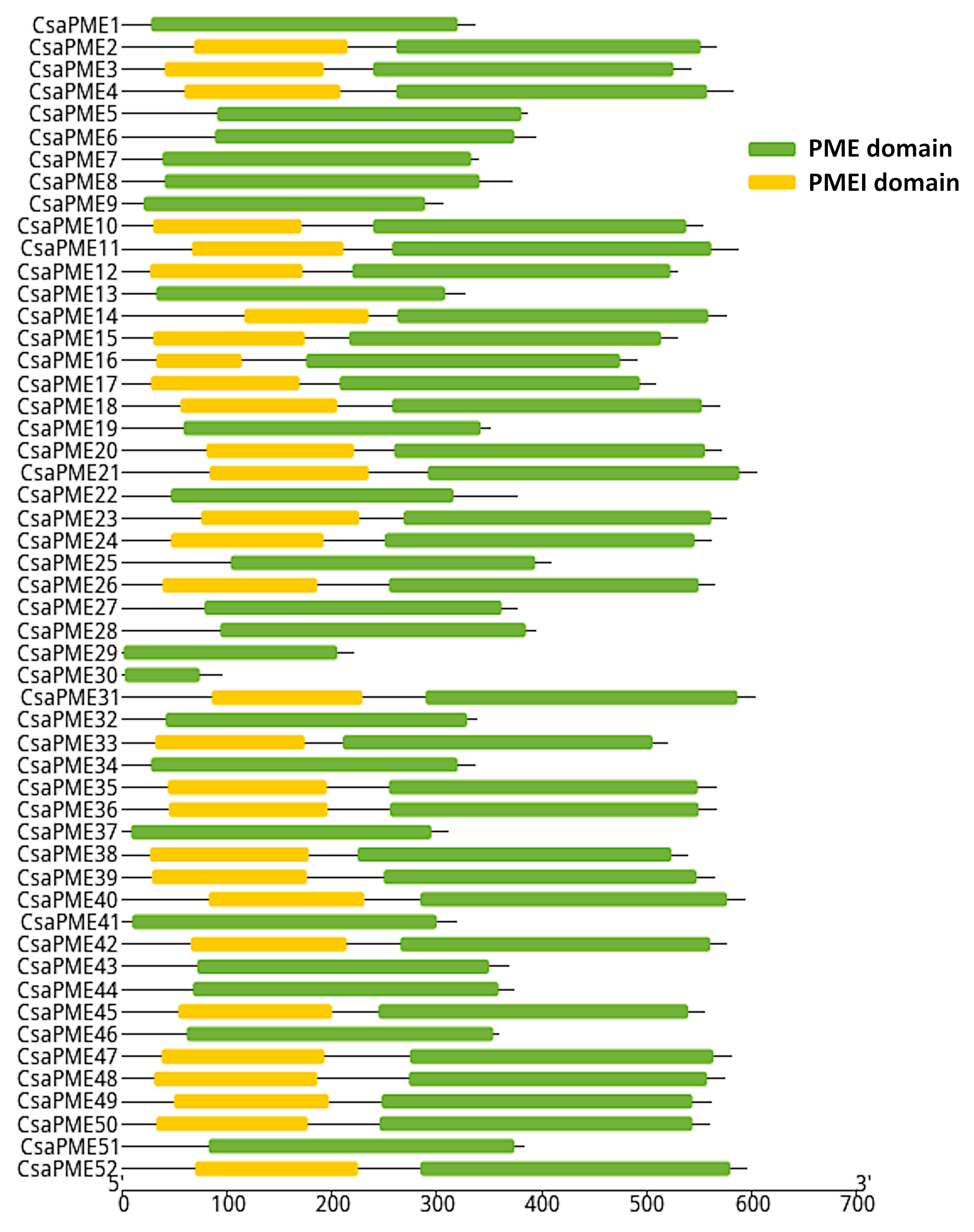

### Figure S6

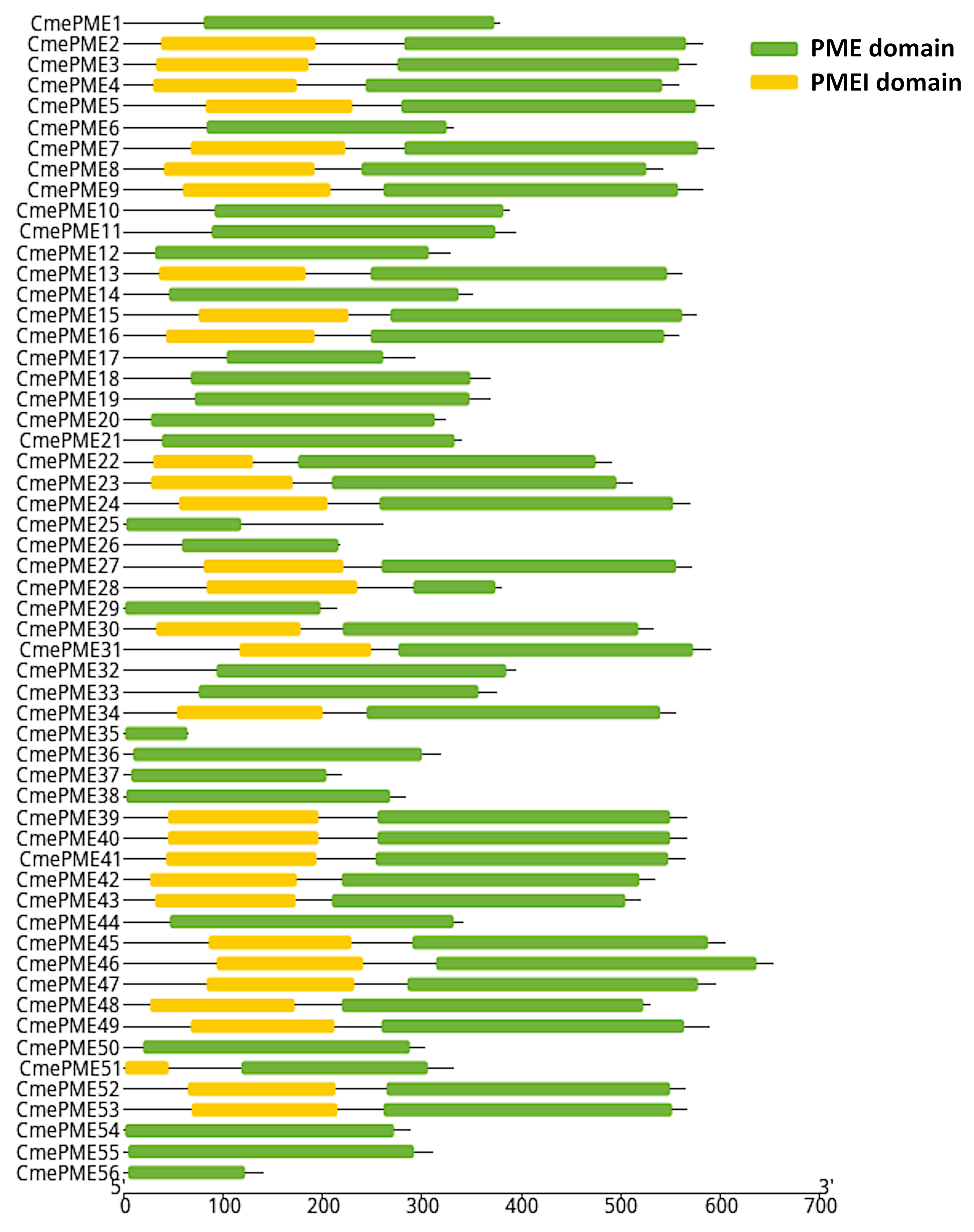

### Figure S7

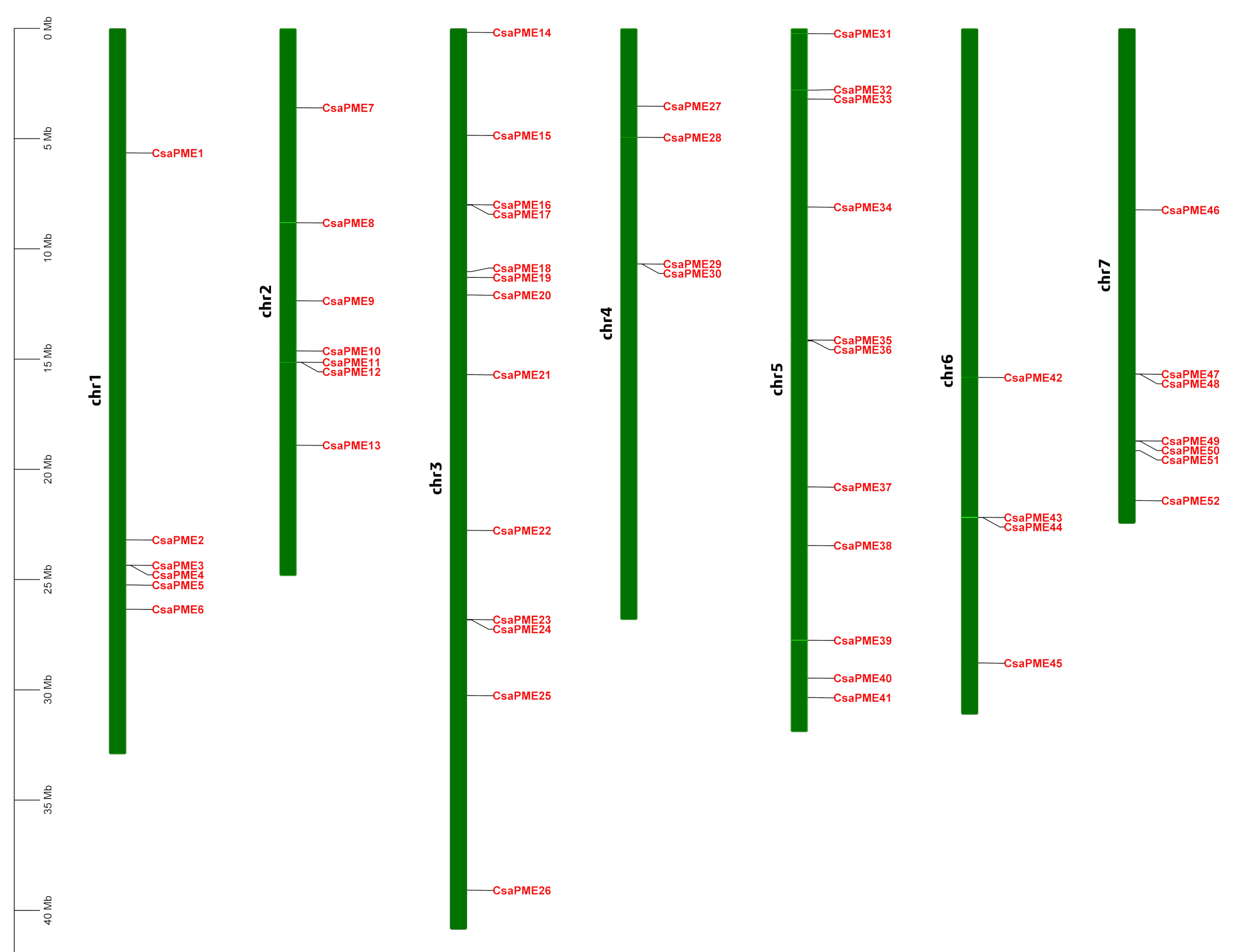

### Figure S8

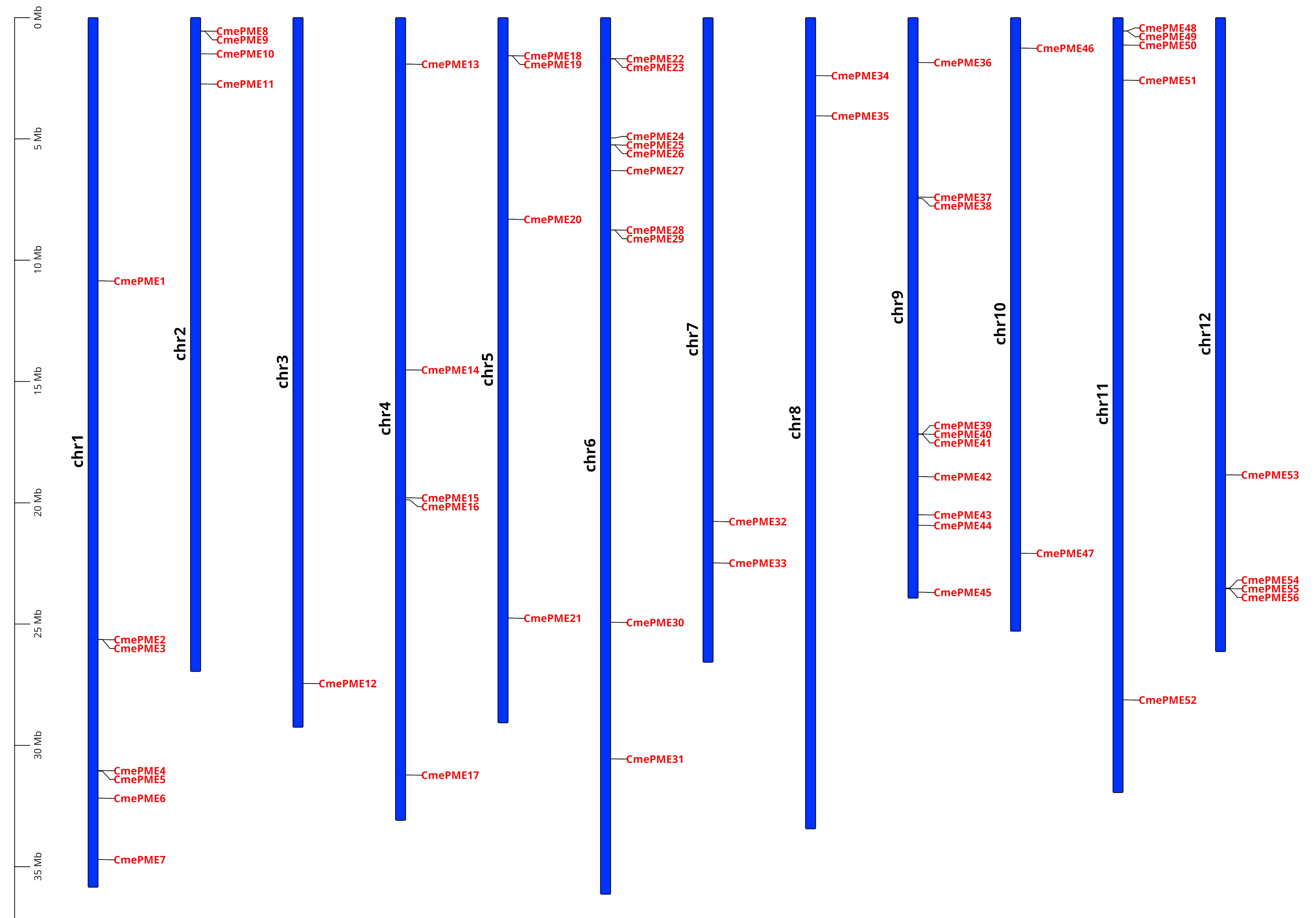

### Figure S9

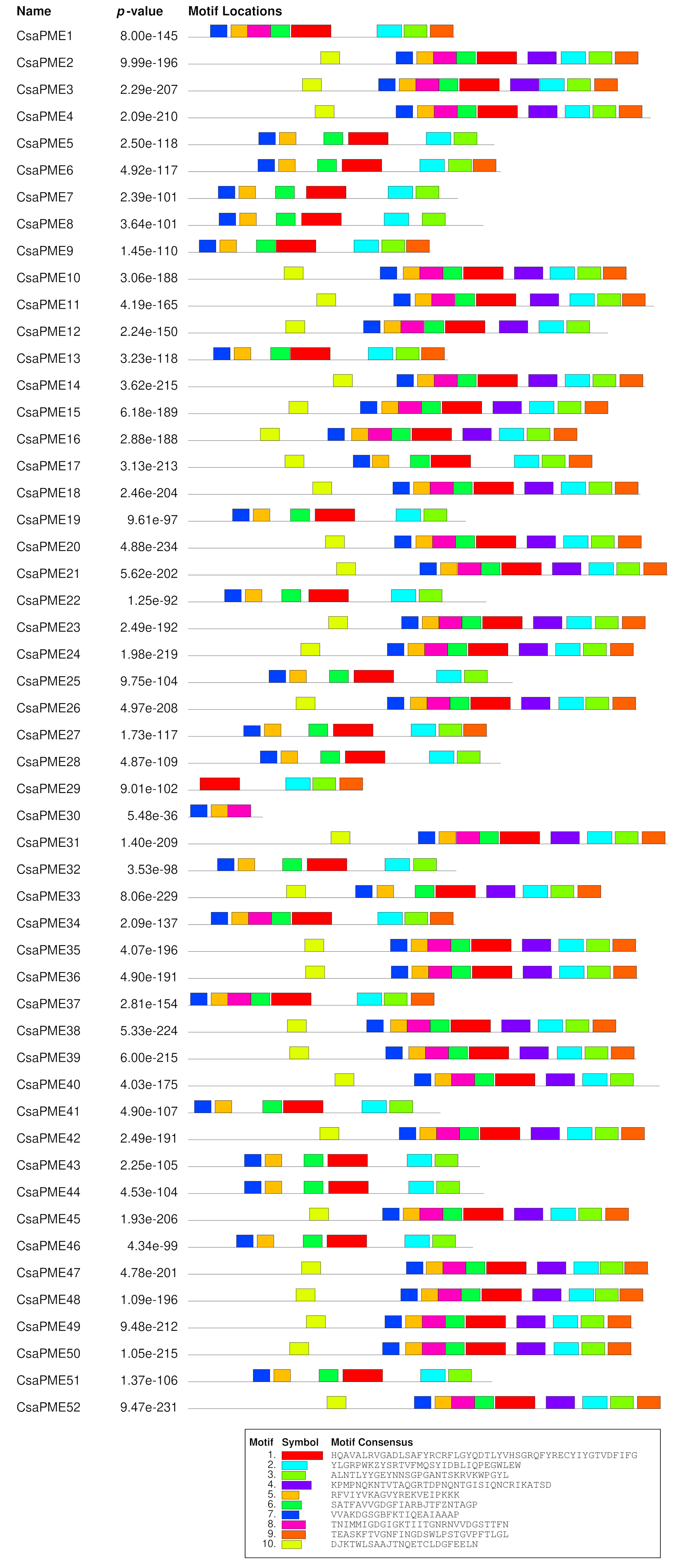

### Figure S10

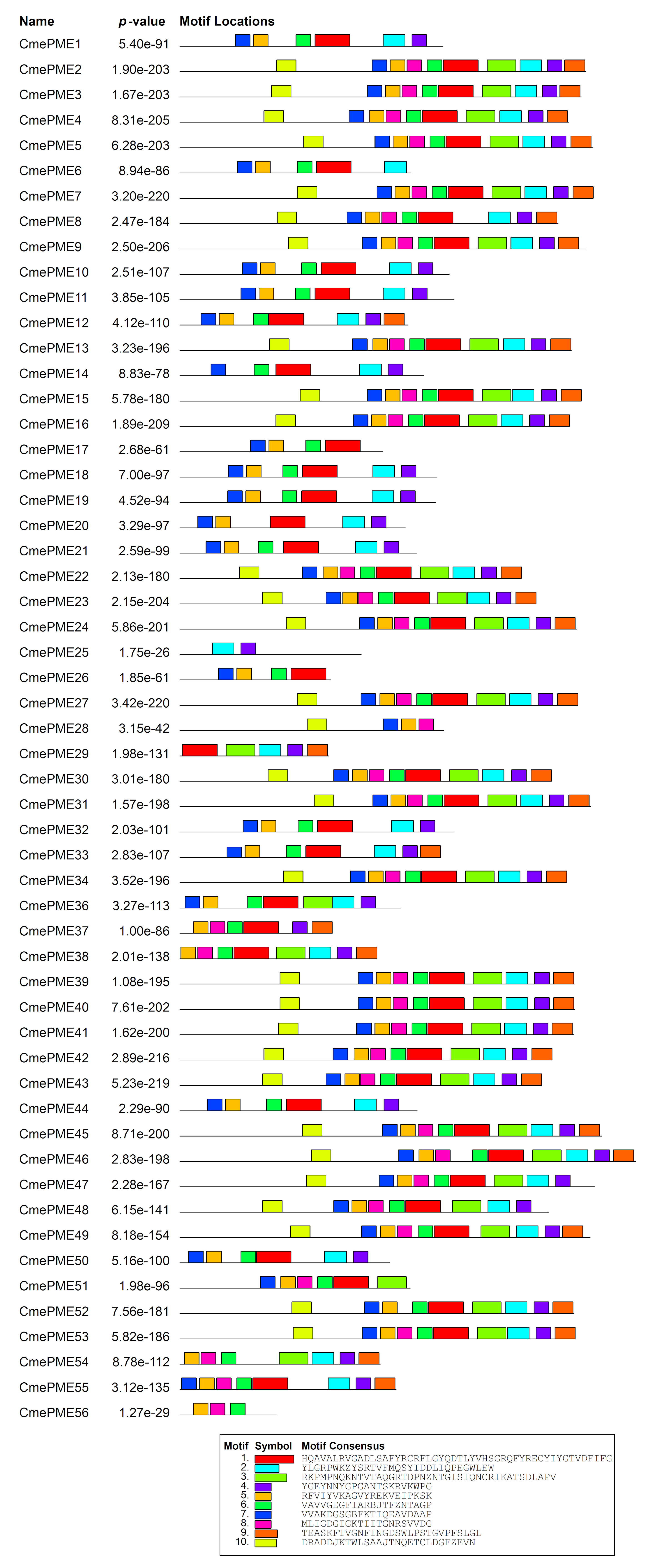

### Figure S11

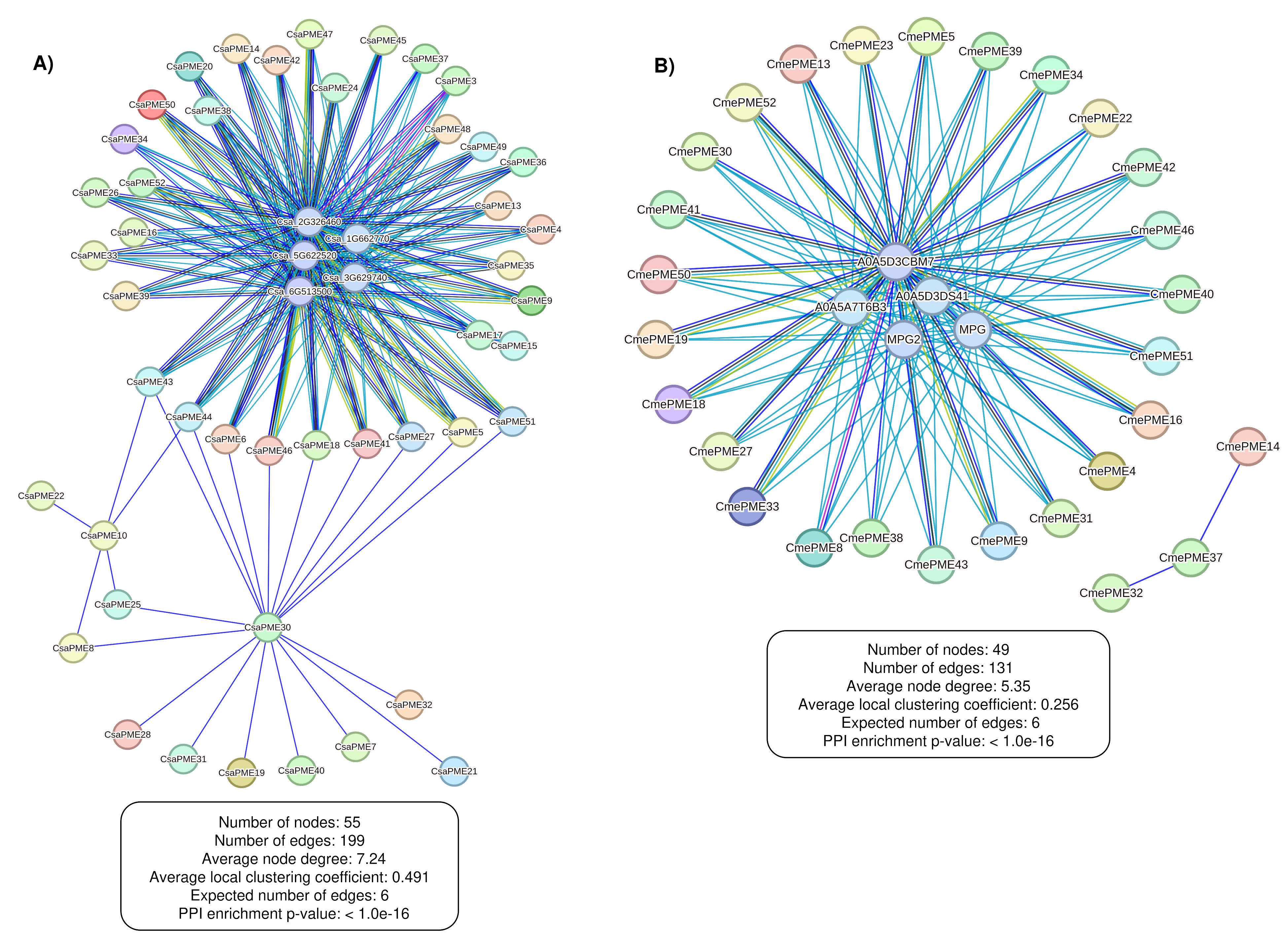

### Figure S12

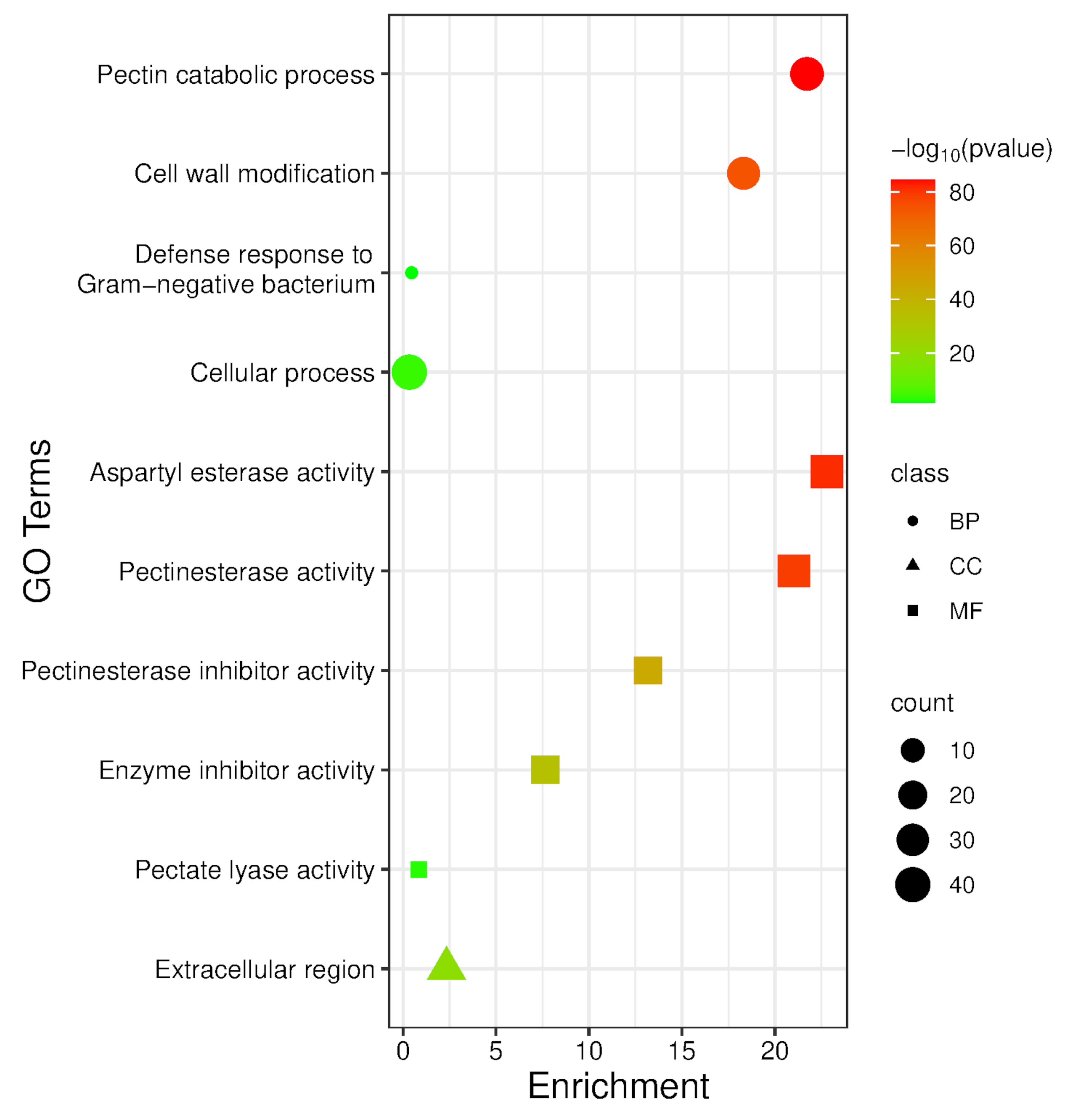

### Figure S13

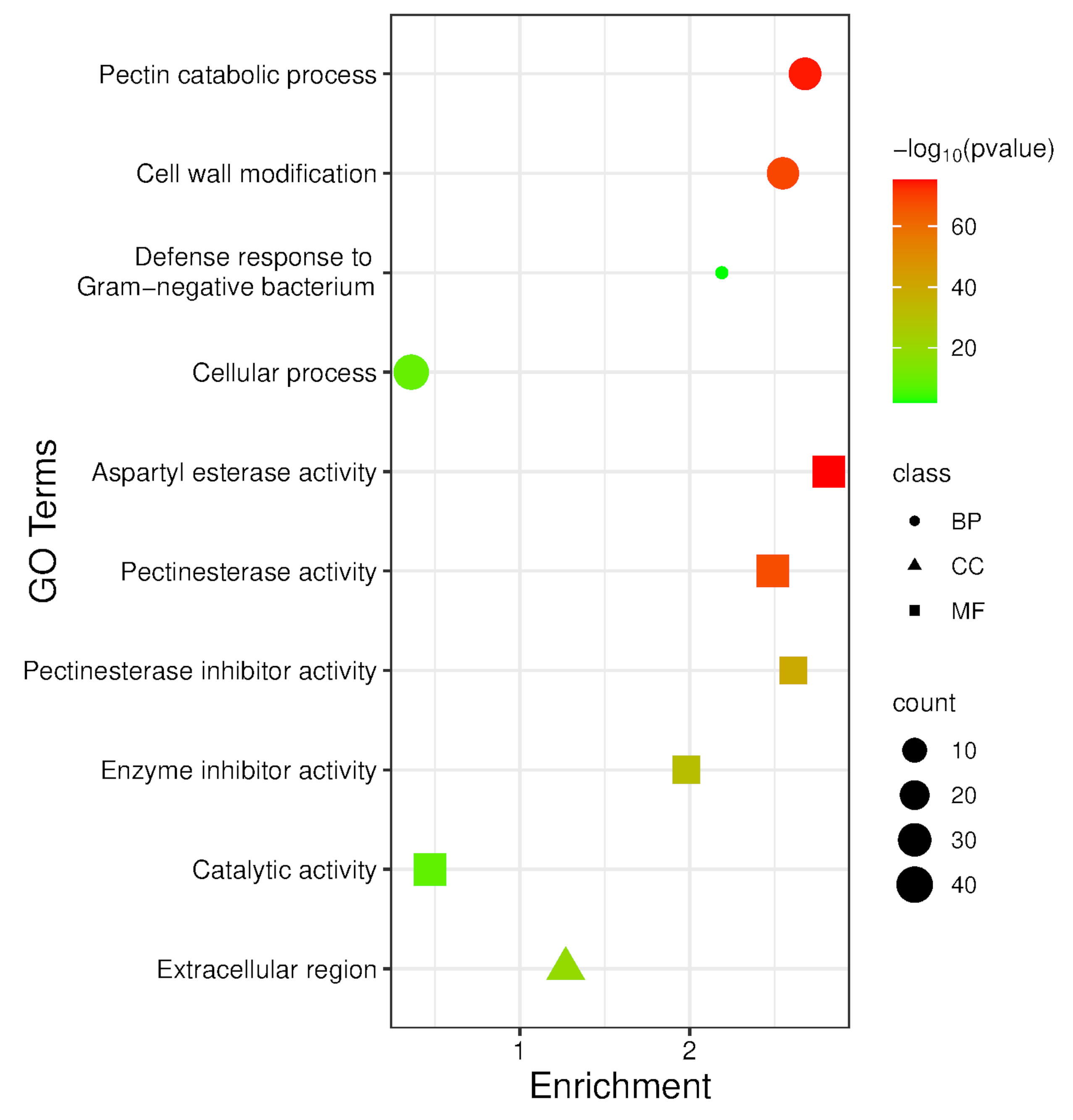

### Figure S14

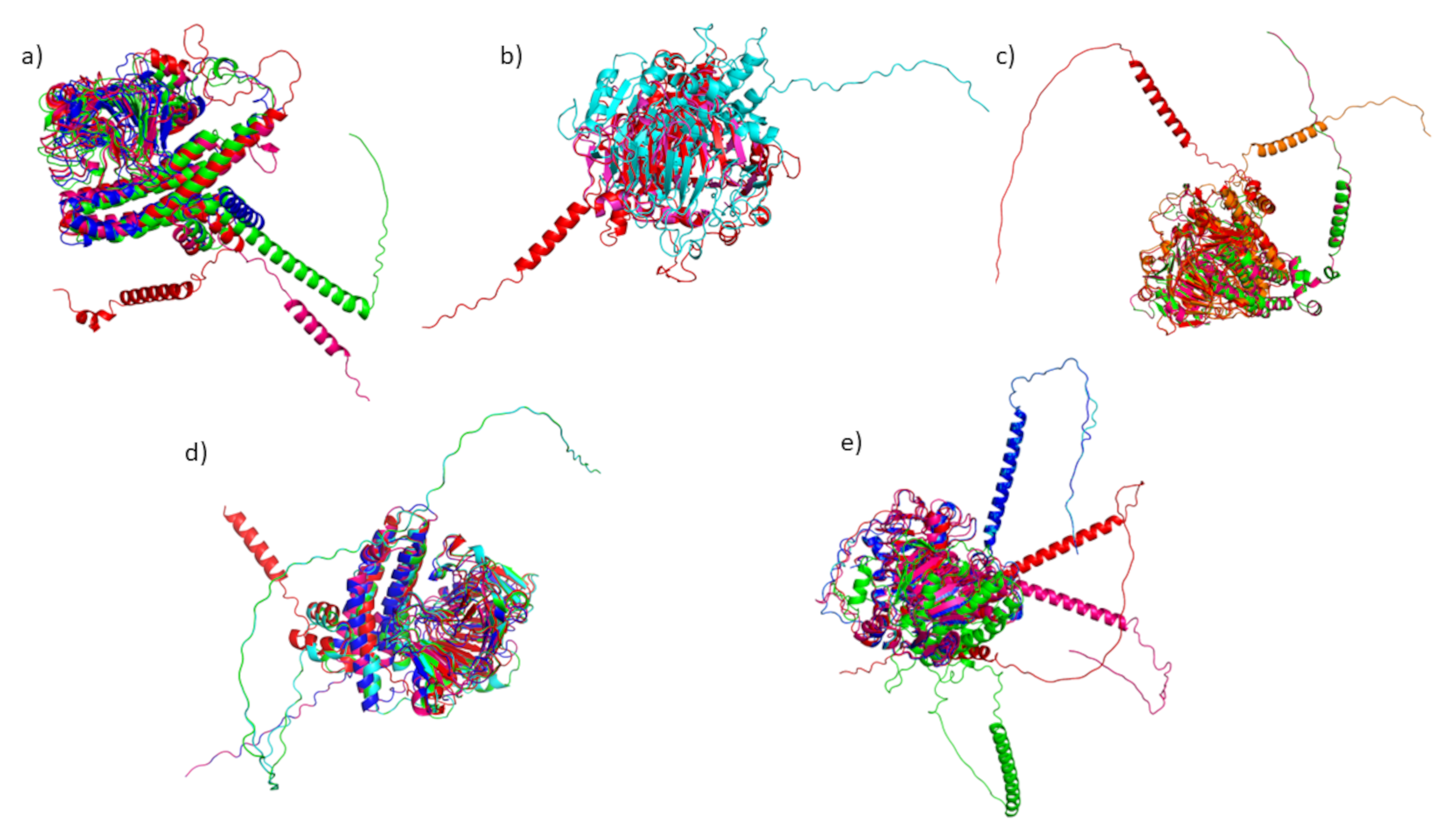
